# BRD4 directs mitotic cell division by inhibiting DNA damage

**DOI:** 10.1101/2023.07.02.547436

**Authors:** Tiyun Wu, Haitong Hou, Anup Dey, Mahesh Bachu, Xiongfong Chen, Jan Wisniewski, Fuki Kudoh, Chao Chen, Sakshi Chauhan, Hua Xiao, Richard Pan, Keiko Ozato

## Abstract

BRD4 binds to acetylated histones to regulate transcription and drive cancer cell proliferation. However, the role of BRD4 in normal cell growth remains to be elucidated. Here we investigated the question by using mouse embryonic fibroblasts with conditional Brd4 knockout (KO). We found that Brd4KO cells grow more slowly than wild type cells: they do not complete replication, fail to achieve mitosis, and exhibit extensive DNA damage throughout all cell cycle stages. BRD4 was required for expression of more than 450 cell cycle genes including genes encoding core histones and centromere/kinetochore proteins that are critical for genome replication and chromosomal segregation. Moreover, we show that many genes controlling R-loop formation and DNA damage response (DDR) require BRD4 for expression. Finally, BRD4 constitutively occupied genes controlling R-loop, DDR and cell cycle progression. We suggest that BRD4 epigenetically marks those genes and serves as a master regulator of normal cell growth.

## INTRODUCTION

BRD4 is a BET family bromodomain protein, expressed in most cells (Dey et al., 2000; Wu and Chiang, 2007). It binds to acetylated histones through the two bromodomains (Dey, 2003; Kanno et al., 2014). BRD4 also binds to P-TEFb, RNA polymerase II, Mediators and others to stimulate transcriptional elongation of numerous genes (Jang et al., 2005; Kanno et al., 2014; Pelish et al., 2015; Whyte et al., 2013). It was reported that BRD4 acts as a kinase and histone acetylase (Devaiah et al., 2016). BRD4 broadly occupies intra and intergenic regions of the genome. It binds on expressed genes with a peak binding at/near the transcription start site (TSS), although it also binds to enhancers, often found distant from the genic region (Devaiah, 2013; Dey et al., 2019; Hnisz et al., 2013; Kanno et al., 2014; Loven et al., 2013; Whyte et al., 2013). BRD4 is essential for embryonic development, and Brd4 knockout (KO) is early embryonic lethal (Nishiyama et al. 2006; Houzelsten et al., 2002). BRD4 facilitates proliferation of many cancer cells (Shi and Vakoc, 2014). Small molecule inhibitors (BETi) arrest cancer growth by blocking interaction of BRD4 with acetylated histones (Filippakopoulos et al., 2010; Loven et al., 2013; Pelish et al., 2015). BETi are thought to be attractive anti-cancer drugs, as they are reported to disrupt oncogene enhancers, and have a minor effect on normal cell growth (Delmore et al., 2011; White et al., 2019; Zuber et al., 2011). More recently developed PROTAC based BRD4 inhibitors are also reported to inhibit cancer growth (Raina et al., 2016). PROTAC-induced BET protein degradation has been used as a therapy for castration-resistant prostate cancer (for ARV-825, Li et al., 2018). However, it is not clear whether BRD4 regulates normal cell growth, although growing evidence points to a pivotal role of BRD4 in cell cycle progression of non-oncogenic cells (Devaiah et al., 2016; Dey et al., 2019; Mochizuki et al., 2008).

DNA replication and mitosis are two fundamental events in cell cycle, which are directed by sequential transcription of more than 1,000 genes (Grant et al., 2013). E2F family of transcription factors are central to G1 to S transition (Bertoli et al., 2013; Grant et al., 2013). FOXM1 is another transcription factor that directs S to G2 transition and G2/M progression (Costa, 2005; Grant et al., 2013; Laoukili et al., 2005). ATR and ATM, previously known to control DNA damage (Blackford and Jackson, 2017), are implicated to function at G2/M stage of cell cycle (Kabeche et al., 2018; Saldivar et al., 2018). It has been shown that ATR is activated at the end of replication and ensures completion of S phase (Saldivar et. al., 2018), and it binds to centromere at mitosis enabling propter chromosomal segregation (Kabeche et al., 2018).

Recent reports on *in vitro* cancer cell models demonstrated that control of DNA damage and repair is an integral part of cell growth regulation, which requires BRD4 (Lam et al., 2020, Edwards et.al 2020). Recently published data from cell-based assays and proteomic interaction network analysis showed that BRD4 acts as anegative regulator of transcription-associated RNA-DNA hybrids (R-loops), since inhibition of BRD4 increased R-loop formation, causing DSBs (Kim et al., 2019). These authors showed that depletion of BRD4 by inhibitors led to DNA damage, growth arrest and cell death. BRD4 was reported to be a multifunctional regulator of chromatin binding that links transcriptional activity and homology-directed repair (Barrows, et al., 2022). Further analyses indicated that this DNA damage was a result of R-loop accumulation. However, the question of whether BRD4 depletion causes R-loop accumulation and DNA damage in non-cancerous cells has remained unknown.

We examined mouse embryonic fibroblast in which Brd4 was conditionally knocked out (KO) to determine the role of BRD4 in cell cycle passage and DNA damage. We found that Brd4KO cells were unable to complete replication and failed to execute mitosis. BRD4 inhibitors led to a similar inhibition in human diploid cells. Moreover, Brd4KO cells incurred extensive DNA damage and increased R-loop formation as evidenced by increased γH2AX foci, comet tails and nuclear S9.6 signals, observed at all stages of cell cycle in Brd4KO cells. Transcriptome analysis found that Brd4KO cells are impaired in expressing many histone genes required for replication and those necessary for mitosis. Further, Brd4KO cells were defective in expressing genes involved in R-loop regulation and DNA damage control, including Top2, Topbp1, Atr, Atm, H2ax and p53. It has been well established that Topoisomerase I (Top1), an enzyme that relaxes DNA supercoiling and prevents R-loop formation in head-on conflicts between replication and transcription and maintains genome integrity (Promonet, et al., 2020). Lastly, ChIP-seq analysis found that BRD4 occupied numerous cell cycle genes as well as genes controlling R-loop formation and DNA damage response throughout the entire cell cycle as detected on the promoter, TSS and gene body. Together our work demonstrates that BRD4 epigenetically marks genes controlling R-loop formation and DNA damage response along with many cell cycle genes and ensures ordered cell cycle events and the maintenance of genome integrity.

## RESULTS

### Brd4 deletion impedes cell cycle progression

To assess the role of BRD4 in cell cycle control, we examined mouse embryonic fibroblasts in which Brd4 was conditionally deleted upon tamoxifen (TMX) treatment (Figure S1A-S1C, Dey et al., 2019). Growth curve analysis showed that Brd4KO cells grew more slowly than WT cells during 12 days of culture, and total cell recovery rate of Brd4KO cells was about 1% of WT cells (Figure 1A). To determine the reason for this growth defect, cells were synchronized at G0 by serum starvation for 72 hours (h), and cell cycle progression was monitored by propidium iodide staining after released by serum addition (Figure 1B). Flow cytometry analysis showed that WT cells, upon exiting G0, evidently proceeded through G1, reaching S at 12h-16h and G2/M at 20h. By 24h, the majority of WT cells finished one cell cycle, and returned to G1 (Figure 1C). In contrast, Brd4KO cells were delayed in S entry (16h), then progressed very slowly, leaving many cells at S phase even at 24h and beyond without reenter G1 (Figure 1C; Figure S1D and S1E). The results indicate that Brd4KO cells are defective in G1/S transition (Mochizuki et al., 2008), but also in S-G2/M passage.

**Figure 1.**
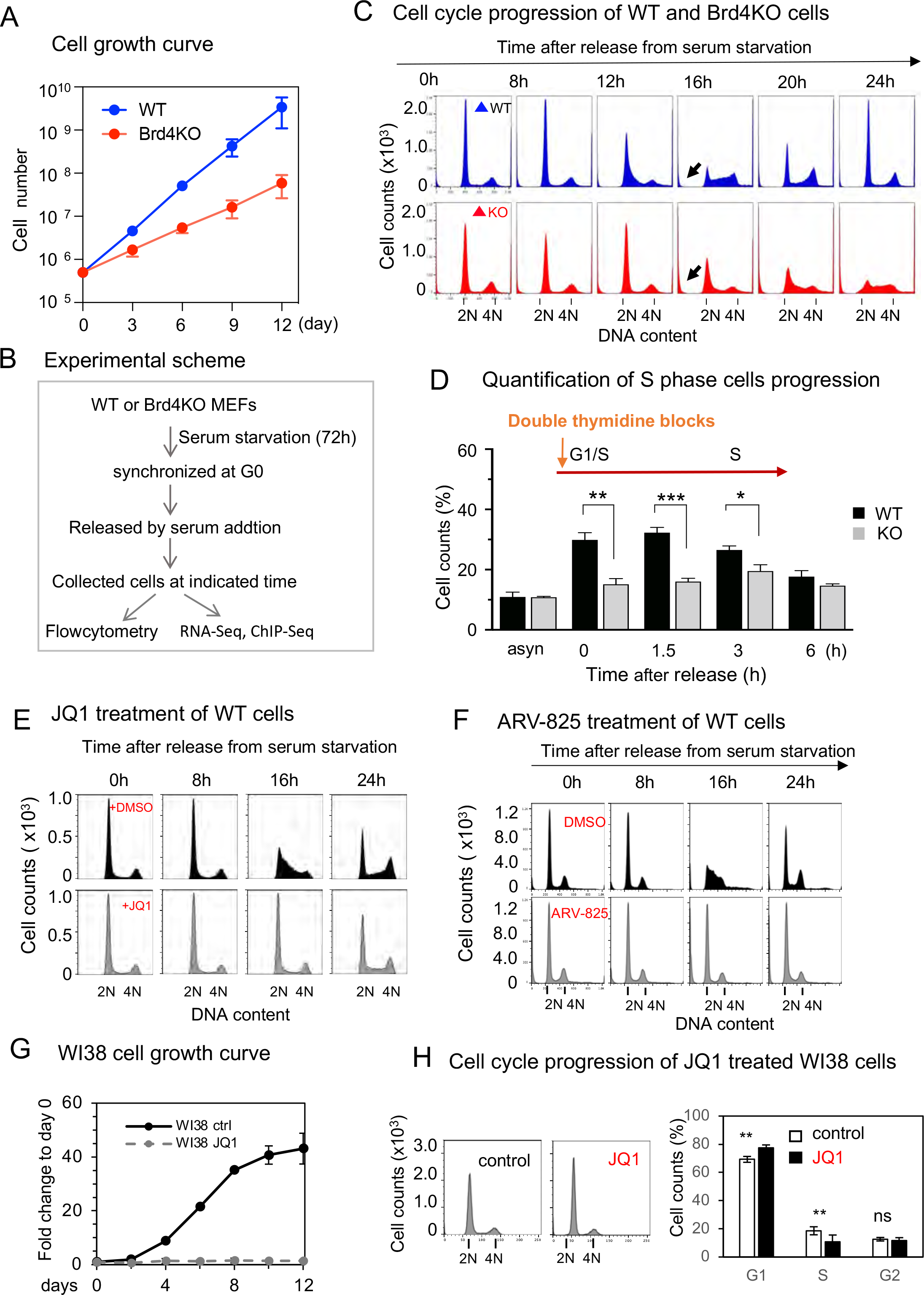
Brd4 deletion and a BET inhibitor impede cell cycle progression. A. Cell growth curve. WT and Brd4KO cells (5 X10^5^) seeded in a 10-ml plate were successively passed, and cell numbers were counted on indicated days. Values are the mean of three biologically separate experiments +/-S.D. B. The experimental scheme. WT and Brd4KO cells were serum starved for 72h, released, and allowed to proceed for 24h and beyond (Figure S1D and E up to 40h). Flowcytometry profiles of PI staining are shown. C. Cells were stained with propidium iodide (25 µg/ml) and DNA contents were determined by flow cytometry. Arrows indicate Sub-2N particles. D. S phase progression after double thymidine block. WT and Brd4KO cells were treated with thymidine (200 µg/ml) twice, released, allowed to proceed up to 6h, and then PI stained cells were analyzed by flowcytometry (Figure S1D). Quantification of S phase cells (the percentage) at each time point is shown. The numbers are the average of three biological repeats. Significance was calculated using unpaired t test (*** P<0.0001; ** P<0.001; * P<0.01). E. WT cells were synchronized by serum starvation for 72h and released. JQ1 (0.5 µg) or DMSO (control) was added at time 0. Flowcytometry profiles of PI stained cells at indicated time are shown. F. WT cells were synchronized by serum starvation for 72h and released as above, and ARV 825 (1.0 µM) or DMSO (control) was added at time 0. Flowcytometry profiles of PI stained cells are shown. G. Growth curve analyses of WI38 cells. WI38 cells treated with 0.5 µM of JQ1 or DMSO, were successively passed, and cell numbers were counted on indicated days. Fold changes are the mean of three biologically separate experiments +/-S.D. H. Cell cycle progression of JQ1 treated WI38 cells. JQ1 or DMSO treated WI 38 cells were stained with PI (25 µg/ml) and DNA contents were determined by flow cytometry (left). Quantification of G1, S and G2 phase cells are shown as the percentage (right). The numbers are the average of three biological repeats. (**p<0.05)

To determine the impact of BRD4 deletion on the progression through S phase, we synchronized cells by thymidine double block at late G1 and early G1/S transition (Choi., et al 2017), and monitored progression with flowcytometry analysis. The counts of WT cells in S phase peaked at 0-3 hours, and G2 cells reached maximum at 6 hours (Figure 1D, S1F). In contrast, S phase progression is significandly blocked (Figure 1D) in Brd4KO cells, as there was very little change in G1 and G2 cell counts from 0h-6h (Figure S1F). These data indicate that Brd4KO cells fail to move through and complete S phase, presumably due to defective replication. Indeed, BrdU incorporation was markedly lower in Brd4KO cells than in WT cells (Figure S1G). Thus, Brd4KO cells were deficient in DNA replication/genome duplication.

We also investigated the effect of JQ1 (a BETi) and ARV-825 (a BRD4 PROTAC inhibitor) on cell cycle progression of WT cells after release from serum starvation (Figure 1E and 1F). JQ1 and PROTAC inhibitor led to the loss of BRD4 (Figure S1K, S1L) and caused clear defects in S phase progression, resulting in a dramatic reduction in S phase cells (Figure S1I and S1J). Brd4KO cell population also contained more sub-2N particles than WT cells (arrows in Figure 1C), indicating increased apoptosis. Indeed, Annexin V staining revealed higher rate of apoptotic cells among Brd4KO cells than among WT cells (Figure S1H, and Figure 6F for additional immunoblotting for apoptosis genes).

To ascertain whether BRD4 is important for the growth of not just mouse cells, we examined the effect of a BETi on WI38 cells, a human primary diploid cell line (Hayflick 1965; Casella et al., 2019). In the presence of JQ1, WI38 cells showed little increase in cell number during 12 days of culture. In contrast, cells grew logarithmically in the absence of JQ1 till reaching a plateau after 8 days (Figure 1G). Flowcytometry analysis of PI distribution showed a reduction in S phase cells after JQ1 treatment (Figure 1H). These data support a general requirement of BRD4 for proliferations of normal noncancerous mouse and human cells.

### BRD4 is essential for proper expression of genes important for cell cycle progression

To identify cell cycle genes regulated by BRD4 in cultured fibroblasts, we performed RNA-seq analysis of WT and Brd4KO cells synchronized by serum starvation (G0) and at various times after release from serum starvation (Figure S2A). In WT cells, there were 2,860 genes showing FC>2 with p-value <0.05 relative to cells at 0h. We designated these genes as cell cycle regulated genes (Figure 2A; Table S1). These genes were then classified into G0, G1, S, G2 and M specific genes according to the time of expression and the published classification (Figure 2B; Heat map and GO analysis in Figure S2B and S2C) (Grant et al., 2013). Supporting our designation, 751 of the 2,860 genes were previously assigned as cell cycle genes in human cells (Figure 2C) (Grant et al., 2013). We found that 2,166 genes were differentially expressed in Brd4KO cells (Figure S2D; Table S2), and that 564 genes were cell cycle genes (Figure 2D). Of these 564 genes, 455 were significantly downregulated in Brd4KO cells, indicating that these genes required BRD4 for proper expression (Figure 2E). These genes were expressed in a stage specific manner, either at G0, G1, S, or G2-M, signifying that BRD4 is required at each cell cycle stage (Figure 2F; Table S3). Many of the S and G2/M genes downregulated in Brd4KO cells are well known for their roles in DNA replication and mitosis (see the list on Figure 2F). For example, genes involved in DNA replication such as Mcms, Orcs, and some E2Fs were downregulated in Brd4KO cells (Figure 2F and S2E). It is worth noting that the expresson of a master proliferation-associated transcription factor, FoxM1 (Grant et al., 2013; Wierstra I., 2013), was lower in Brd4KO cells (Figure S2B and S2E). G2 and M genes which were downregulated in Brd4KO cells also included cyclins and kinases that signal entry into and completion of mitosis, e.g., Cdc20, Cdc25b, Cdk1, Cdk2, Ccnb1, Aurka, b, and Plk1 (Figure 2F, S2E).

**Figure 2.**
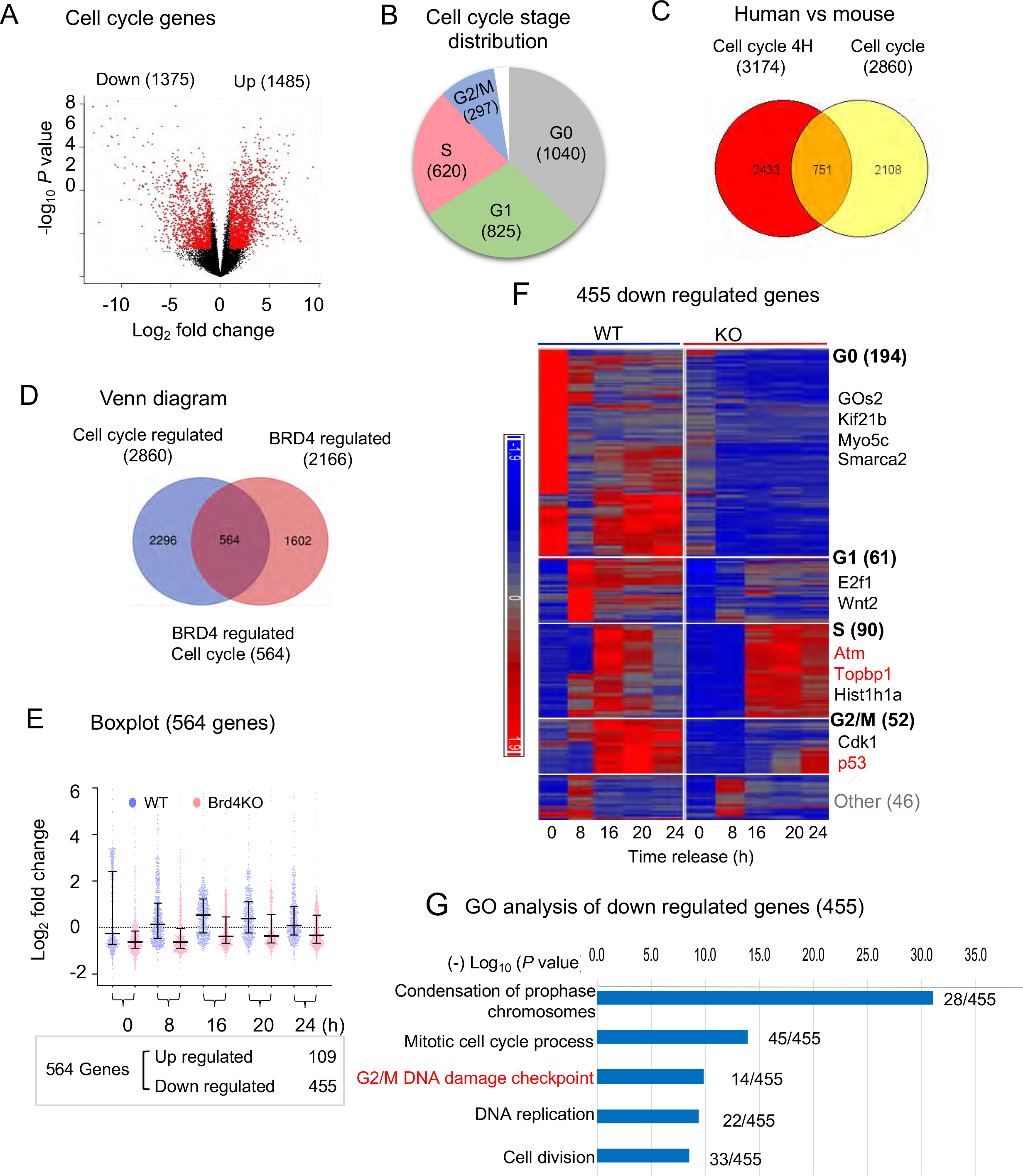
RNA-seq identification of BRD4 dependent cell cycle genes. A. Volcano plot identifying cell cycle regulated genes in WT cells. The Y axis represents -log10 *P* Values and the X axis fold changes (log2 fold change). 2,860 genes (FC > 2, *p* value <0.05) between 0, 4, 8, 12,16, 20 and 24h were designated as cell cycle regulated genes. Peak expression of each gene was assigned based on peak correlation to the idealized phase expression profiles ((Figure S 2b). B. Pie chart showing the number of cell cycle genes expressed at G0, G1, S and G2/M. C. Venn diagram showing the relationship between cell cycle-regulated genes (3669) in four human cancer cell lines (in U2OS, HeLa cells, Foreskin fibroblasts and HaCaT cells)^20^ and cell cycle regulated genes in mouse (2860). D. Venn diagram showing the relation between cell cycle regulated genes and BRD4 regulated genes. 564 genes (in the overlapped area) represent cell cycle regulated genes that are also regulated by BRD4. E. Boxplots representing expression ranges of each gene in WT (Blue) and Brd4KO cells (red) at each time point with *p* value < 0.0001 by two-way ANOVA analysis. Of 564 BRD4 regulated genes, 455 genes were reduced in Brd4KO cells, while 109 genes were increased in expression. F. Heatmap representation of Brd4 dependent cell cycle genes. Genes are grouped according to the cell cycle stages. The number in parenthesis indicates the number of genes downregulated in Brd4KO cells for each stage. Names of representative cell cycle genes are shown. G. GO analysis of BRD4 dependent cell cycle genes. GO terms showing the highest p-values are shown. The number of the genes identified in this study vs total gene numbers in the category are shown on the right.

### Brd4KO reduces histone expression at S phase and hampers induction of G2/M master regulators

RNA-seq analysis showed that transcription of as many as 52 of the 86 core histone genes was downregulated in Brd4KO cells (Heat map in Figure S2E). These genes are mapped to the core histone gene clusters 1 and 2, transcribed along with DNA replication during S phase (Marzluff et al., 2002). The striking reduction in histone gene transcription in Brd4KO cells (Figure S2E) prompted us to examine histone protein levels in synchronized Brd4KO cells. Histones were acid extracted and subjected to SDS PAGE analysis (Figure 3A). Quantification showed that the amount of H2A, H2B/H3, and H4 increased about 2-fold in WT cells after S phase as would be expected, but little or no increase was observed for any of the histones in Brd4KO cells (Figure 3B). The reduction of core histone expression may lead to cell cycle defects. Furthermore, genes encoding centromere/kinectcore proteins CenpA, CenpE, CenpN and CenpF and Kinesin family proteins (Kifs, Walzak, et al., 2010) (Figure 2F, 2E and S2E) were also downregulated in Brd4KO cells (Figure 3C). The RNA-seq data were further confirmed by qRT-PCR (Figure S3A), and protein levels were determined by western blotting analysis (Figure S3B). Thus, consistent with an obligatory doubling of core histones and centromere/kinectcore components for successful genome duplication, we observed the reduction in BrdU incorporation in Brd4KO cells (Figure S1G), which correlated to the reduced expression of core histones.

**Figure 3.**
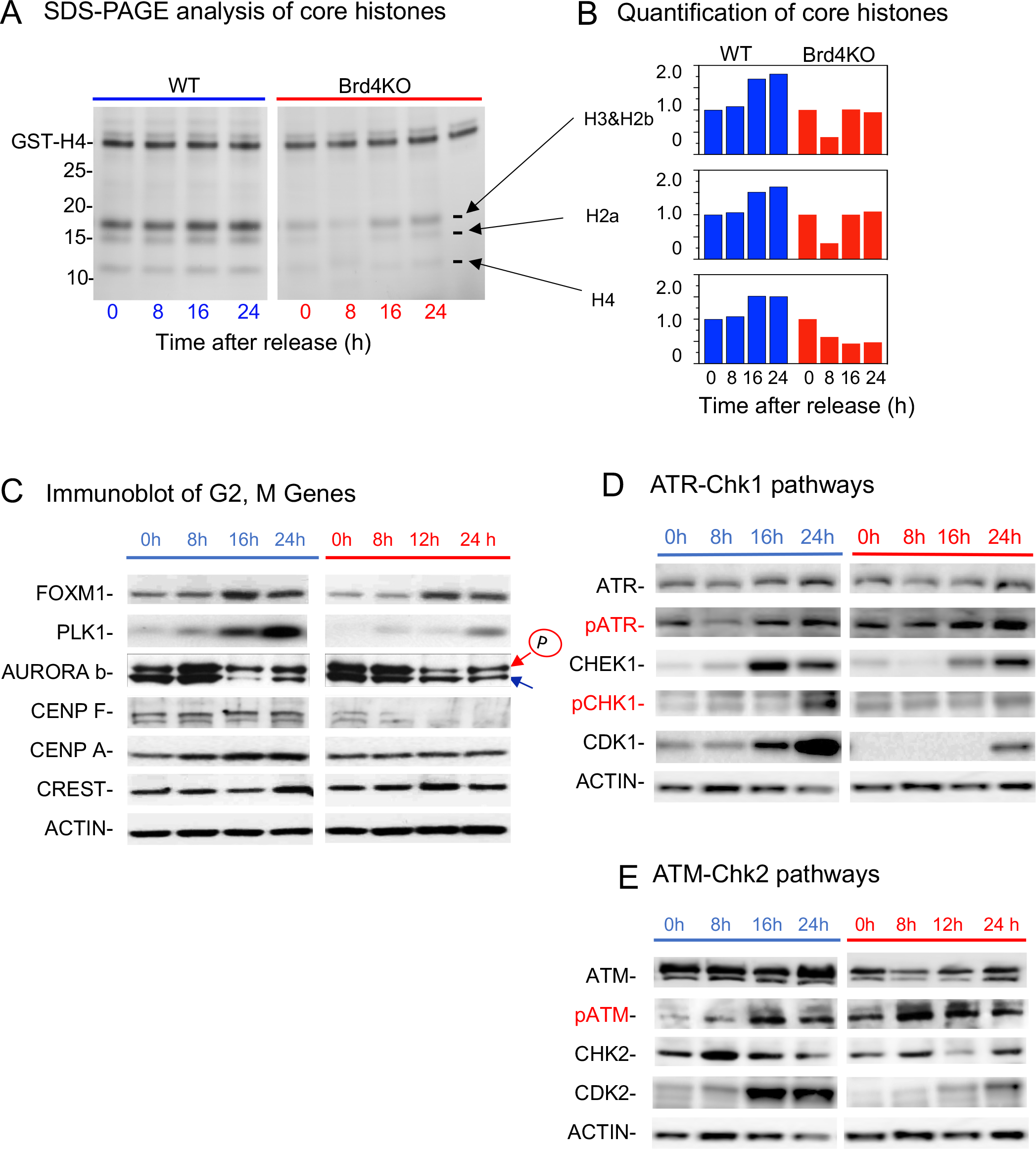
Protein levels for Histones, FOXM1 and ATR/ATM are reduced in Brd4KO cells. A. Acid extracted histone (2µg) preparations were separated on 12% NuPAGE gel and stained with SYPRO Orange. GST-H4 was added in histone extracts to be used as loading control. B. Core histones were normalized against GST-H4, quantified by the ChemiDoc MP Imaging System. C, D, E. Immunoblot analyses of proteins for ATM-CHK2 pathway, ATR-CHK1 pathway and G2/M passage in synchronized WT and Brd4KO cells. βACTIN was used as a loading control. In E, antibody for BRD4 was tested to confirm its absence in KO cells (Figure S3b).

We also confirmed protein levels of major G2/M genes such as FOXM1, ATR and ATM, since their transcript levels were reduced in Brd4KO cells. As shown in Figure 3C, the amount of FOXM1 was lower in Brd4KO cells throughout the cell cycle (Figure S3B). In addition, the levels of PLK1, CENP-F, and CENP-A, downstream targets of FOXM1, required for ordered mitotic processes were lower in Brd4KO cells than in WT cells. As shown in Immunoblot data in Figure 3D and 3E, the amounts of ATR and ATM, and the downstream kinases, CHK1, CHK2, CDK1 and CDK2 were also lower in Brd4KO cells, consistent with the lower transcript levels. We noted that despite reduced protein expression of ATR, the amount of phosphorylated ATR was higher in Brd4KO cells than WT cells, however, we did not observe a clear difference in phospho-ATM levels (Figure 3E, S3B). These results provide the first suggestive evidence for DNA damage in Brd4KO cells (Figure 3D, S3B). In conclusion, BRD4 is essential for proper expression of genes important for genome replication and chromosome segregation.

### BRD4 constitutively marks genes regulating cell cycle and genome stability

BRD4 has been shown to distribute broadly over the mouse and human genome, and within the genic regions including promters, exons and introns, and the occupancy correlates with active gene expression (Hargreaves, 2009; Nicodeme, 2010; Zhang, 2012; Dey, 2019). Here we performed ChIP-seq analysis of synchronized cells to examine genome-wide BRD4 distribution at each stage of the cell cycle. The global BRD4 binding increased when exiting G0 and entering G1 (8h), and peaks at S (16h) followed by a slight decline at G2/M (20h (Figure 4A).).

**Figure 4.**
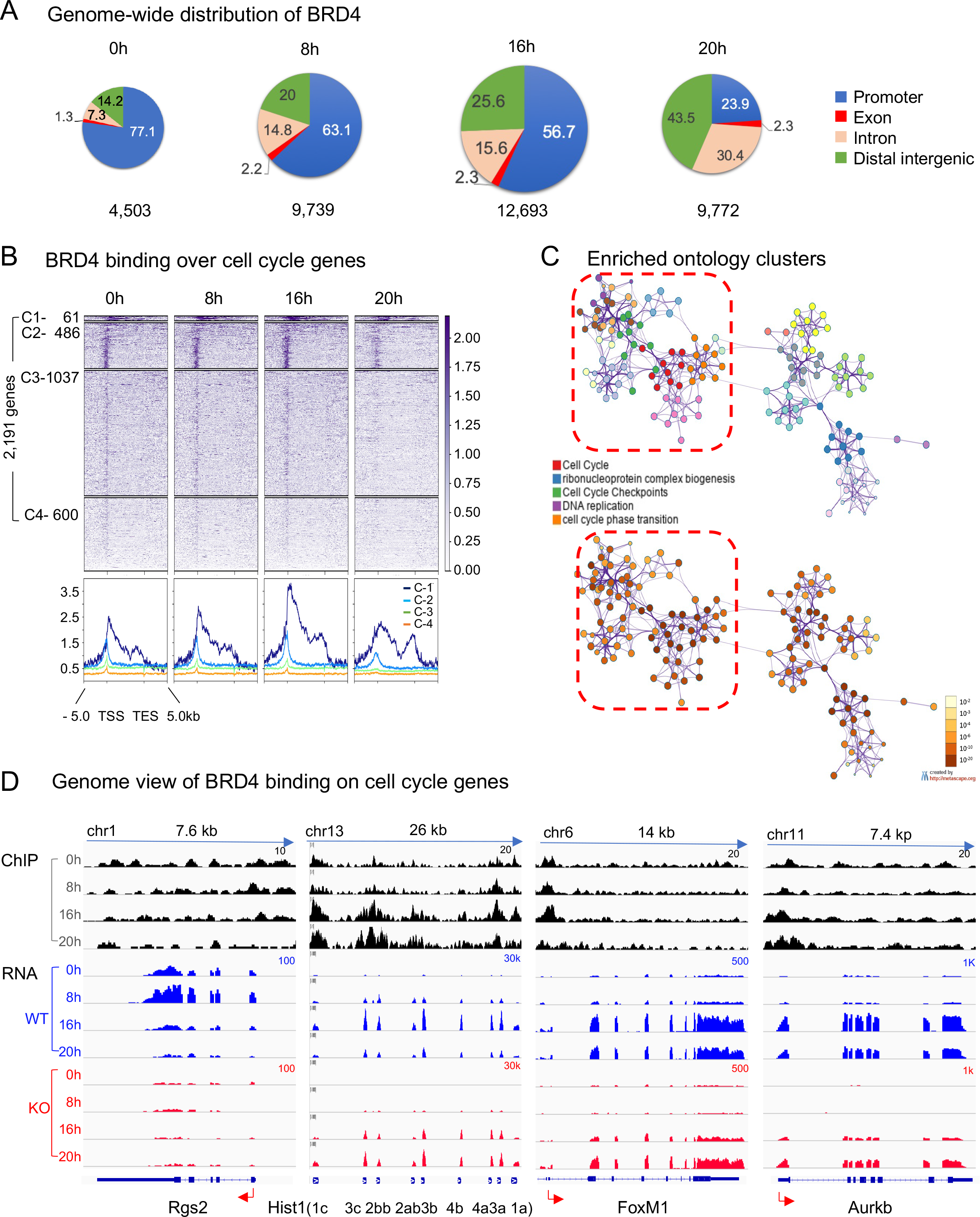
BRD4 constitutively marks cell cycle genes with the peak occupancy at S phase. A. Pie charts depicting genome-wide distribution of BRD4 binding at G0, G1, S and G2/M (0h, 8h, 16h, 20h). The numbers below indicate the total peak counts at each stage. BRD4 distribution over the promoter (blue), genic (pink+ red) and distal intergenic regions (green) are shown. Note the highest peak counts at S (16h). B. Top, heat maps, BRD4 binding on cell cycle regulated genes are plotted over the TSS, gene body, TES +/-5Kb at indicated cell cycle stage. Genes are aligned according to BRD4 signal intensity (high to low), with Cluster 1 (C1) showing the highest BRD4 signals, while C4 the lowest. The numbers of genes belonging to each cluster are shown on the left. Bottom, average BRD4 distributions. Note the highest BRD4 signals at S phase. C. Metascape based pathway analysis for Cluster1 to Cluster 3 genes (Figure 4B). The network of enriched terms is color-coded by cluster ID, where nodes that share the same cluster ID are typically close to each other (top) and colored by p-value, where terms containing more genes tend to have a more significant p-value (Bottom). D. IGV profiles of BRD4 occupancy (top) and RNA-seq peaks (middle and bottom) for select cell cycle genes in WT and Brd4KO cells. Gene names and the exon-intron organization are shown below.

We then examined BRD4 occupancy on 2,191 cell cycle genes as defined by RNA-seq data. The heat maps in Figure 4B depict BRD4 occupancy over the promoter, transcription start site (TSS), gene body and the transcription end site (TES) and the flanking 5Kb regions. As shown in Figure 4B, cell cycle genes occupied by BRD4 were aligned according to BRD4 signal intensity, from the highest to the lowest (from top to bottom). These genes were divided into four clusters (C1 to C4). C1, with the highest BRD4 signals, contained 61 genes. C2, C3 and C4 contained 486, 1037 and 600 genes, respectively (see Table S4 for gene lists in each cluster). The average BRD4 binding profile for each cluster is shown at the lower panel of Figure 4B. In each cluster, BRD4 binding was the highest at the promoter/TSS region, and trailed over the gene body, similar to the binding profile seen with other genes in other cells (Dey et al., 2019; Loven et al., 2013). Although BRD4 bound to both cell cycle and non-cell cycle genes, binding intensity was significantly higher on cell cycle genes than on non-cell cycle genes (Figure S4A). It was remarkable that the pattern of BRD4 binding was similar throughout all cell cycle stages, also seen when peak BRD4 binding was plotted at the center (Figure S4B). These results indicate that BRD4 binds to cell cycle genes at all stages, even though the expression of these genes is stage specific. Metascape-based GO analysis of genes in C1 through C3 revealed enrichment with the terms such as cell cycle, cell cycle transition, and DNA replication (Figure 4C). To verify constitutive BRD4 occupancy on cell cycle genes, we plotted BRD4 binding on G0, G1, S and G2/M specific genes identified in Figure 2F separately and observed again constant BRD4 binding to these genes throughout cell cycle (Figure S4C). Interestingly, S phase genes displayed the highest BRD4 intensity, while G0 genes were the lowest. Genome views of genes showing constitutive BRD4 binding but stage specific RNA expression are presented in Figure 4D and S4D. Genes representing G1 (Rgs2, Ddit4), S (7 histone genes, Chek1) and S-G2 (FoxM1, Aurkb, Cdk1, Ccnb1) are shown for BRD4 binding (Top) and RNA expression in WT (Middle, blue) and in Brd4KO cells (Bottom, Red). These genes were always bound by BRD4, although binding intensity varied somewhat at different stages. Taken together, BRD4 binds numerous cell cycle genes throughout the entire cell cycle, but directs stage specific transcription. We later found that BRD4 also constitutively occupied genes that regulate R-loop formation and DNA damage and was required for their RNA expression (described below, Figure 6H, and S6G).

### Brd4KO cells are defective in mitotic entry and undergo mitotic failure

The above genome-wide analyses showed that BRD4 is critical for proper regulation of S and G2/M genes. To further investigate cell cycle progression of Brd4KO cells, we performed live cell imaging of cells expressing a GFP-tagged histone H3.1. Images of randomly growing population revealed many healthy mitotic cells in WT culture (Figure 5A). In contrast, cells with large nuclei, presumably at late S phase dominated Brd4KO culture, with very few mitotic cells. Some of the Brd4KO cells accompanied small, micronucleus-like structures (Figure 5A) (Crasta et al., 2012; Lewis and Golsteyn, 2016). To further ascertain whether Brd4KO cells are unable to pass through G2/M, we performed nocodazole block experiments. Nocodazole inhibits microtubule polymerization and arrests cells at prometaphase where chromosomes begin to line up at the mitotic plate (Nishiyama et al., 2012). Asynchronously growing WT and Brd4KO cells were treated with nocodazole for 8h, and prometaphase arrested cells were identified by DNA stain (arrows in Figure 5B). Whereas a large fraction of WT cells was arrested by Nocodazole, very few Brd4KO cells displayed mitotic arrest (Figure 5B, on the right). Flowcytometry profiles showed that nocodazole treatment led to accumulation of 4N cells in WT cells, whereas many Brd4KO cells remained in mid to late S, indicating the inability of Brd4KO cells to reach prometaphase (Figure S5A). Consistent with these results, other mitotic inhibitors, (+)-S-trityl-L-cysteine (STLC) and MG132, did not bring Brd4KO cells to 4N (Santaguida and Amon, 2015; Skoufias et al., 2006) (Figure S5A and S5B). To further corroborate defective mitotic entry, we immuno-stained nocodazole treated WT and Brd4KO cells with antibody for H3pS10, a marker of chromosome condensation and mitosis (Wei et al., 1999). Brd4KO or ARV-825 treated WT cell populations contained fewer H3pS10-positve cells with condensed chromosomes (Figure 5C at bottom). Immunoblot analyses of cells synchronized by serum starvation showed a marked increase in H3pS10 signals in WT cells after 16 hours consistent with the timing of mitotic entry, whereas H3pS10 signals remained low at all stages in Brd4KO cells (Figure 5D).

**Figure 5.**
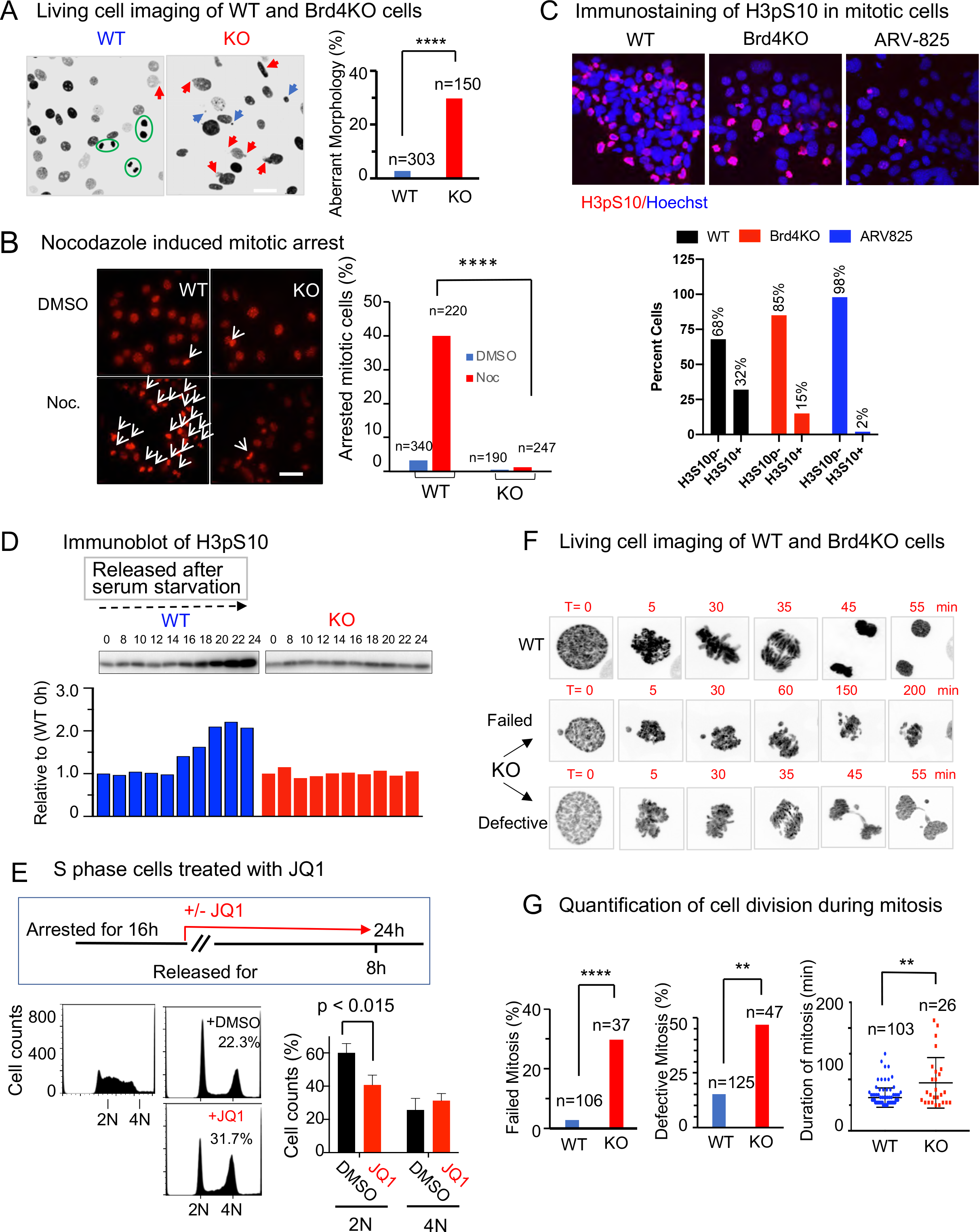
Live cell imaging reveals catastrophic mitotic failure with chromosomal mis-segregation in Brd4KO cells. A. Representative snapshots of live WT and Brd4KO cells expressing H3.1-GFP. Arrows Indicate large, morphologically unusual cells. Scale bar = 50 μm. Percentages of cells with unusual morphology were quantified from 9 separate fields from WT and Brd4KO cells, respectively (left), quantification of cells with aberrant morphology. N indicates the number of cells counted (right). B. Nocodazole induced mitotic arrest. Asynchronous WT and Brd4KO cells were treated with Nocodazole (100 ng/ml) or DMSO (control) for 8h, stained with Hoechst 33342, and viewed on confocal microscopy. Scale bar=10μm. Mitotically arrested cells are marked by arrow (left). Quantification of mitotically arrested cells (%). N indicates the number of cells counted (right). C. Immunostaining of H3pS10 in mitotic cells. Nocodazole (50ng/ml) arrested WT, Brd4KO cells and ARV-825 treated WT cells, were immunostained with H3pS10 antibody. Mitotic specific chromatins were indicated with H3pS10 staining WT (left), Brd4KO (middle) and ARV-825 treated WT (right) cells (Pink), and counterstained with Hoechst 33342 (blue) for DNA (top). Quantification of mitotically arrested cells (% of H3pS10-or H3pS10+ cells). (bottom). D. Immunoblot of H3pS10. WT and BRD4KO cells were harvested at indicated time points after serum starvation and release. Acid extracted histones were separated on 12% NuPAGE gel and blotted with H3S10ph antibody (top). H3S10ph bands were normalized against the loading control (High molecular weight band in the acidic extract) and subsequently quantified using Image J software (bottom). E. S phase cells treated with JQ1. WT cells were synchronized by serum starvation, released and allowed to proceed for 16 h. JQ1 (0.5µM) or DMSO was added at this time and cells were further incubated for 8h (top). Cells were stained with propidium iodide and DNA contents were analyzed by flow cytometry (right). The percentages of cells at G1(2N), and G2/M (4N) are quantified (left). F. Live cell imaging of WT and Brd4KO cells. Frames of representative films monitoring mitosis. Note that most WT cells (Top panel) completed mitosis within 55 min. Some Brd4KO cells failed to achieve mitotic cell division and disintegrated (middle panel). Other Brd4KO cells were delayed in metaphase-anaphase progression and displayed chromosomal mis-segregation (bottom panel). G. Quantification of failed (left), defective as bridge during mitosis progression (middle) and duration of mitosis (right) and (Figure S5D) in Brd4KO and WT cells.

We next tested JQ1 for its effect on G2/M passage in WT cells. Cells were synchronized by serum starvation, and JQ1 was added at 16h when cells were at S phase (Figure 5E, top). FACS analysis showed that majority of cells without JQ1 treatment completed one cell cycle and were back in G1 by 24h (Figure 5E, Bottom). In contrast, JQ1 treated cells remained at or before G2/M stage. These results show that JQ1 interferes with S to G2/M passage in normal cells, independently of its inhibition on G1/S transition. Thus, Brd4 is important for proper cell cycle progression at every stage of the cell cycle.

It should be noted that although most Brd4KO cells encountered delay in mitotic entry, some cells managed to reach mitosis. We monitored mitotic progression by time lapse imaging of WT and Brd4KO cells undergoing mitosis (Figure S5C video and snapshots 5F and S5D). As shown in Figure 5F top and S5D top, WT cells completed mitosis within 55 minutes, starting from chromosomal condensation, alignment on the mitotic plate, sister chromatid separation to the formation of two daughter cells. In contrast, few Brd4KO cells showed signs of mitosis. Although some Brd4KO cells exhibited chromosomal condensation, many of them failed to line up on the mitotic plate (Figure 5F, middle and bottom; Figure S5D bottom). Consequently, these cells did not progress beyond this stage and eventually disintegrated presumably by apoptosis. Some other Brd4KO cells reached a metaphase-like state, followed by chromosomal segregation, but taking much longer time and showing a higher frequency of chromosomal miss-segregation, such as lagging chromosomes and chromosomal bridges (Figures 5F and S5B bottom), similar to previously reported anaphase DNA bridges (Liu et al., 2014). Quantification in Figure 5G showed a much higher frequency of failed mitosis and defective segregation of Brd4KO cells than WT cells. These data indicate that BRD4 deficiency can lead not only growth arrest, but also a loss of genome integrity.

### BRD4KO cells incur DNA damage

Taken together, our results highlight the importance of BRD4 in DNA replication and mitotic cell division, thus defects in cell cycle progression of BRD4KO cells are possibly a result of DNA damage (Floyd et al., 2013; Donati 2018; Zhang, et al., 2018; Barrows et al. 2022). To assess DNA damage, WT and Brd4KO cells were immunostained with antibody against phosphorylated H2AX (уH2AX) (Bonner, et al., 2008). As shown in Figure 6A and S6A, most Brd4KO cells displayed intense γH2AX foci over the nuclei, while WT cells had many fewer γH2AX positive nuclei. Moreover, consistent with the conserved function of BRD4, JQ1 treatment of human WI38 cells resulted in significantly higher γH2AX signals compared to control cells treated with vehicle alone (Figure 6B).

**Figure 6.**
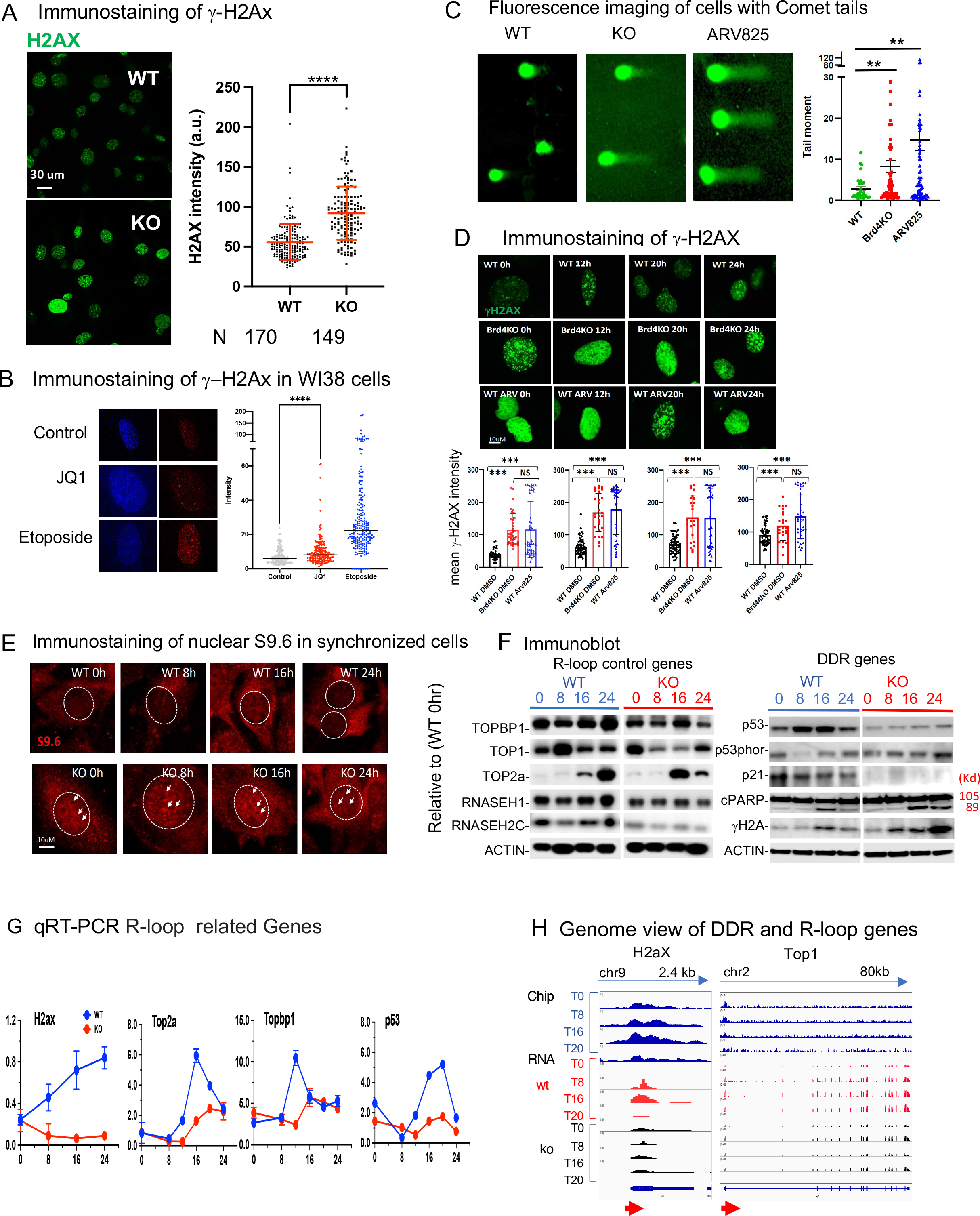
R-loop accumulation and DNA damage following BRD4 depletion. A. Immunostaining of γH2AX in randomly growing cells. WT and Brd4KO cells were immunostained with γH2AX antibody (left). Quantification of γH2AX Fluorescence (right), Significance was assessed using two-tailed unpaired t test. B. Immunostaining of γH2AX in JQ1 treated WI38 cells . WI38 cells were treated with JQ1 (2.5µM) for 24h and immunostained with γH2AX antibody (left). IF intensity in each cell was quantified (right). Significance was assessed using Mann-Whitney test (**** P<0.0001). (right), Comet assay. WT, Brd4KO and ARV-825 treated WT cells were subjected to Comet assay in alkaline conditions. Comet tails were quantified using tail moment of open comet program (n = 3 independent experiments (right). Significance was calculated using unpaired t test. C. Immunostaining of γH2AX in synchronized cells. WT and Brd4KO cells were synchronized by serum starvation, released and incubated for indicated times. WT cells (top), Brd4KO cells (middle) and WT cells were treated with ARV-825 (low) were immunostained with γH2AX antibody at indicated times. Quantification of mean γH2AX intensity was performed using two-tailed unpaired t test (bottom. ***P < 0.001). D. Immunostaining of nuclear S9.6 in synchronized cells. WT and Brd4KO cells synchronized by serum starvation and allowed to proceed to indicated times after release were immunostained with S9.6 monoclonal antibody and antibody to Nucleolin (to mask signals from nascent nucleolar RNA). Representative fluorescence images of nuclear S9.6 (white cycles) are shown. Quantification of nuclear S9.6 intensity was performed using ImageJ and is shown in Figure S6B (*P <0.05 to ***P < 0.001). E. Immunoblot detection of R-loop and DDR ol genes. WT and Brd4KO cells synchronized by serum starvation, released and allowed to proceed for indicated times. Nuclear extracts (20 <u>µg</u>) were separated on 4-12% NuPAGE gel and blotted with antibodies indicated on the left. Left and right panels represent antibodies for R-loop and DDR proteins, respectively. See quantification in Figure S6F. F. qRT-PCR analyses of R-loop and DDR genes. Transcript expression of H2ax, Top2a, Topbp1 and p53 in WT and Brd4KO cells synchronized as in Figure 6F were tested by qRT-PCR. Blue line for WT and red line for Brd4KO cells. (See Figure S6E for Top1, Top3b, and p21). G. IGV profiles of BRD4 occupancy (top) and RNA-seq peaks (middle and bottom) for H2ax and Top1 genes in WT and Brd4KO cells. Gene names and the exon-intron organization are shown on the top and botto, respectively. (See IGV profiles for Rnaseh1, topbp1, Top2b, top3b, p53 and p21 in Figure S6G).

To further verify that BRD4 depletion leads to DNA damage, we performed Comet assays that detect cells with DNA damage (Olive and Banath, 2006). This is a single cell gel electrophoresis assay where cells with DNA strand breaks produce comet tails which are visualized by microscopy. As shown in Figure 6C, many Brd4KO cells but not WT cells exhibited discernible comet tails. As expected, WT cells treated with ARV825 which degraded BRD4 also produced comet tails (Figure 6C).

To determine whether the extent of DNA damage varies during cell cycle, we examined γH2AX signals in synchronized WT, Brd4KO, and ARV825 treated WT cells (Figure 6D).. Cells released from serum starvation were cultured for 0h, 12h, 20h, 24h, and then immunostained for γH2AX, as these samples were enriched with G0, G1, S and G2/M, respectively. As shown in Figure 6D, Brd4KO cells and ARV825 treated cells showed much higher γH2AX signals than vehicle treated cells at all stages. Overall, levels of γH2AX signals were always similar in Brd4KO and ARV825 treated cells, and consistently higher than in WT cells (bottom). In addition, flow cytometry was carried out to detect γH2AX in synchronized WT and Brd4KO cells. FACS data in Figure S6B showed that γH2AX signals were higher in Brd4KO cells than WT cells at all times (at bottom).

We then examined the expression of genes for DNA damage recognition/repair (DDR) pathway. The tumor suppressor p53 when phosphorylated, directs DNA repair, cell cycle and apoptosis (Williams and Schumacher, 2016; Engeland K., 2022). Although the levels of total p53 were reduced in Brd4KO cells (Figures 6F and S6F), the amount of phosphorlated p53 was a little higher in Brd4KO cells than in WT cells, indicating activation of DDR in Brd4KO cells, likely due to activation of ATR (Figures 3D and S3B). These data are in line with the increased γH2AX levels in Brd4KO cells (Figure 6F right). However, p21, activated by phospho-p53, was lower in Brd4KO cells than in WT cells (Figures 6F right panel and S6F), indicating impaired DDR in Brd4KO cells. Further suggesting the complexity of DDR defects in Brd4KO cells, PARP1 cleavage products were higher in Brd4KO cells than in WT cells (Figures 6F and S6F) (Mashimo et al. 2021; Ray Chaudhuri and Nussenzweig, 2017).

### Brd4KO cells accumulate R-loops

It has recently been reported that BRD4 depletion in some cancer cells results in accumulation of R-loop, which further leads to DNA damage (Lam et al., 2020; Edwards et al., 2020; Kim et al., 2019). R-loop, an RNA-DNA hybrid, is produced during transcription and post-transcriptionally, and can be visualized by immunostaining with the monoclonal antibody S9.6 specific for RNA-DNA hybrid (Bou-Nader et al., 2022). To assess whether Brd4KO cells accumulate R-loop and at what stage of the cell cycle, synchronized WT and Brd4KO cells were stained with the S9.6 antibody and an anti-nucleolin antibody, the latter to mask nascent RNA (Sollier et al., 2014). As shown in Figure 6E, we detected distinctly higher intranuclear S9.6 signals in Brd4KO nuclei compared to WT nuclei (Figure S6C). As expected, S9.6 staining was found in the cytoplasm of all cells, likely representing mitochondrial R-loop (Sollier et al., 2014). Furthermore, overlapping γH2AX-S9.6 double staining signals suggest physical proximity of R-loop and DNA damage (Figure S6D, enlarged image at the bottom, quantification on the right). These results reinforce the idea that R-loop accumulates in cells lacking Brd4, which triggers DDR.

Next, we performed immunoblotting analysis to determine if factors controlling R-loop formation depend on BRD4 for proper expression. As shown in Figure 6F (left), TOP1, TOP2a, TOPBP1, RNAseH1 and RNAseH2C1 were reduced in Brd4KO cells relative to WT cells throughout cell cycle (Figure S6F).

We then examined the possibility that BRD4 is required for transcription of factors critical for R-loop control and DDR. qRT-PCR data from synchronized cells showed that transcript levels of H2ax, Top2a, Topbp1, p53, as well as Top1, Top3b, RnaseH2b and p21 were much lower in Brd4 KO cells than in WT cells (Figure 6G and S6E). Except for H2ax which was expressed throughout cell cycle, other genes were expressed after S and G2/M. These results were consistent with RNA-seq data, in that IGV profiles of transcript peaks revealed that they were lower in Brd4KO cells (two lower panels in Figure 6H and S6G). Thus, Brd4KO cells are deficient in factors controlling R-loop formation and invoke defective DDR.

Furthermore, these genes were occupied by BRD4 throughout all stages of cell cycle as shown in the IGV profiles of ChIP-seq peaks (top panel, Figure 6H, and S6G). BRD4 binding peaks were found not only at the promoter/TSS area, but over the gene body, similar to the binding patterns of many cell cycle genes (Figure 4D, Figure S4D). Thus, BRD4 epigenetically marks R-loop and DDR genes and directs their transcription. These results highlight a close similarity to cell cycle genes (a Model in Figure S6H). Together, R-loop and DDR genes require BRD4 for expression and that the absence of BRD4 leads to impaired R-loop resolution, dysregulated DDR along with defective replication and mitosis.

## DISCUSSION

Our results that Brd4 drives cell cycle progression in normal mouse cells and human diploid cells is a notable observation, given that BRD4 has been reported to selectively promote cancer cell growth (Zubar et al., 2011; Loven et al., 2013). Analyses of synchronized WT and Brd4KO cells by serum starvation, thymidine double block at S phase, mitotic block by nocodazole revealed that BRD4 directs S phase passage independent of G1/S regulation; the absence of BRD4 impeded completion of S phase, and passage to G2/M. Consistent with the importance of BRD4 in this step, frequency of mitotic cells was markedly lower in Brd4KO culture than in WT cell population. Moreover, the few Brd4KO cells that reached mitotic stage were either disintegrated or produced daughter cells with unequal chromosomal segregation. Thus, BRD4 has a decisive role in two fundamental events in cell growth, i.e., genome replication and mitosis (Grant, et al., 2013). Together, BRD4 directs not only G1/S transition as noted before (Mochizuki et al., 2008), but the progression in the later stages of cell cycle, which has not been fully appreciated before.

Transcriptome analysis revealed that more than 450 genes involved in cell cycle progression require BRD4 for expression, the numbers far higher than other cell cycle regulators such as E2Fs and FOXM1 (Grant et al., 2013). Many genes necessary for DNA replication/genome duplication and mitosis were dependent on BRD4. Remarkably, BRD4 is required for transcription of 52 of the 86 histone genes including all core and linker histones, which were downregulated in Brd4KO cells. Given that BRD4 was bound to individual histone genes shown by our ChIP-seq data, BRD4 likely drives transcription of each histone gene. Histone gene regulation in mammalian cells has not been fully deciphered, although some factors including YY1 and NPAT are shown to control histone expression (Marzluff et al., 2002; Nizami et al., 2010; Ye et al., 2003). Our study adds BRD4 as a critical factor for global regulation of histone gene transcription. In line with RNA levels, histone proteins were severely reduced in Brd4KO cells, verifying that BRD4 is essential for histone production. In addition, some factors that direct DNA replication, such as members of the Mcm and Orc families were downregulated in Brd4KO cells, illustrating that BRD4 is critical for chromosomal duplication. It is likely that coordination of DNA replication and subsequent chromatin assembly is grossly defective in Brd4KO cells. Moreover, BRD4 was required for expression of many key proteins necessary for mitosis, those involved in the formation of kinetochores and centromere, and regulation of spindle fibers, including Cenps and Kifs plus various G2/M kinases. Thus, transcriptome data are consistent with phenotypic defects most prominent in Brd4KO cells undergoing mitosis.

One of key features of our study is that BRD4 occupied the target cell cycle genes at all times, despite that their transcription was restricted to specific stages. BRD4 binding was observed even at G0, and the binding intensity fluctuated during cell cycle and tended to be higher at the time of transcription. In general, the highest BRD4 occupancy was seen at the proximal promoter/TSS area, but its binding was evident on the gene body as well. In light of previous reports that BRD4 plays a role in gene marking, while it directs transcription of target genes (Dey et al., 2009; Behera et al., 2019), it is interesting that BRD4 epigenetically marks cell cycle genes on one hand, and directs their stage specific transcription on the other (See a Model in Figure S6H).

The data that ATR was activated in Brd4KO cells led to an unexpected revelation that BRD4 is involved in DNA damage control. Brd4KO cells exhibited many more γH2A foci and comet tails than WT cells. That WI38 cells when treated with JQ1 also led to DNA damage lends further credence to the role of BRD4 in preventing DNA damage. Our results are in agreement with earlier reports that BRD4 inhibitors induce DNA damage and growth arrest in cancer cells (Lam et al., 2020; Kim et al.,2019; Edwards et al., 2020). Moreover, we showed that a series of genes required for DNA damage response (DDR) are markedly down regulated in Brd4KO cells, including Atr and Atm, H2ax, p53 and its target p21, leading to reduced protein expression. It is clear that BRD4 plays a major role in directing DDR gene transcription. Another key observation is that these DDR genes were bound by BRD4 throughout cell cycle, the same feature as that noted for cell cycle genes.

Recent reports showed that increased R-loop formation accounts for DNA damage in BRD4 depleted cancer cells, providing mechanistic insight into the basis of DNA damage (Lam et al., 2020; Kim et al., 2019; Edwards et al., 2020). R-loop, a RNA-DNA hybrid, found in about 5% of mammalian genome has a fundamental role in transcription, replication and telomere maintenance (Niehrs, C and Luke, B 2020; Hegazy, Y et al., 2020, Sollier, et al. 2015). R-loop is produced during transcription and at post-transcriptional steps R-loop also accumulates during replication due to collision of replication fork and transcription. Indeed, Lam et al., (2020) and Kim et al (2019) reported that BRD4 depletion increases R-loop formation likely due to the replication-transcription conflict. In this study, we also observed R-loop accumulation in Brd4KO cells and BRD4 inhibitor-treated WT cells, as evidenced by S9.6 nuclear stain. Notably, however, the levels of S9.6 nuclear signals were similar throughout cell cycle, not restricted to S phase. Thus, R-loop formation in Brd4KO cells is likely attributed to occur through transcription and posttranscriptional steps. This view is consistent with the fact that BRD4 is a part of the transcription elongation complex and helps drive nascent mRNA synthesis (Patel, et al., 2013; Devaiah et al., 2013; Sarai, et al., 2013). Strikingly, a number of genes shown to restrict R-loop accumulation, such as Topbp1, Top2 and Rnaseh1 were downregulated in Brd4KO cells, at RNA and protein levels analogous to DDR genes. Further highlighting the similarity with DDR and cell cycle genes, BRD4 dependent R-loop controlling genes were bound by BRD4 at all times during cell cycle, again with the highest binding at the promoter/TSS sites.

In conclusion, BRD4 orchestrates cell cycle progression by integrating mechanisms of regulating R-loop formation and DNA damage responses (Model in Figure S6H). Together BRD4 ensures genome integrity and healthy cell renewal.

## STAR Methods

### Cells, synchronization, and drug treatments

Embryonic fibroblasts were prepared from Brd4^f/f^ ER^2-^Cre mice and cultured in Dulbecco’s modified Eagle’s medium (DMEM) supplemented with 10% (v/v) fetal bovine serum (FBS) and 1% penicillin-streptomycin (Invitrogen) (Dey et al., 2019). To delete Brd4, cells were treated with 2 µM 4-hydroxytamoxifen (TMX, Sigma) for 96 h. Cells were treated with JQ1 1(Sigma), BET bromodomain PROTAC ARV-825 (Selleckchem.com) Thymidine (Sigma-Aldrich), Nocodazole (Sigma-Aldrich), MG132 (Sigma-Aldrich), and (+)-S-Trityl-L-cysteine (HTLC) (Sigma-Aldrich) at the concentration and duration indicated in Figure Legends.

The WI38 cells were cultured by EMEM with 10%FBS, HEPES, NEAA. Growth Curve for WI-38 primary cells, 50,000 WI-38 cells were plated onto a 12-well plate. After 24 hours, the cells were treated with 0.5 µM JQ1 and then fixed with 4% paraformaldehyde. For staining, the plates were incubated with 0.1% crystal violet in PBS for 30 minutes at room temperature, followed by three washes with water. The dye was dissolved in 1 mL of 10% acetic acid after the plates were completely dried. Cell growth was measured by the absorbance at 590 nm.

For synchronization, WT and Brd4KO cells were incubated in DMEM with 0. 05% FBS for 72h, released in the complete media at indicated times. Cells were fixed with 70% Ethanol, stained with propidium iodide (25 μg/ml) and analyzed by flow cytometry on FACS Caliber interfaced with the Cell Quest software (BD Biosciences).

For thymidine double block used to synchronize cells in late G1/early S (Galgano and Schildkraut, 2006), WT and Brd4KO cells were first treated with thymidine (25 mg/ml) at 37°C for 22h, released in complete media at 37°C for 6h, followed by a second treatment with thymidine (25 mg/ml) at 37°C for 16h, and finally released in complete media, and cells were collected at indicated time points. Cells were then fixed with 70% Ethanol, stained with propidium iodide (25 μg/ml), and analyzed by flow cytometry on BD FACSCalibur interfaced with the Cell Quest software (BD Biosciences).

To detect apoptosis, cells were stained with propidium Iodide and annexin V-fluorescein isothiocyanate (BD Biosciences Pharmingen) according to the manufacturer’s instructions.

### Quantitative real time (qRT)-PCR and RNA-Seq analysis

Total RNA was prepared by Quick-RNA Miniprep Kit (ZYMO research). Gene-specific cDNA was synthesized with random hexamers (Thermo Scientific). Quantitative Real Time-PCR (qRT-PCR) was performed using Fast SYBR^TM^ Green Master Mix (Applied Biosystems, Thermo Fisher Scientific).

For RNA-seq Libraries were prepared from total RNA as above using a Mondrian SP (NuGEN Technologies Inc.) and the Ovation SP Ultralow Library system (NuGEN). Fragments ranging from 250 to 450 bp were subjected to paired-end sequencing on a HiSeq 2000 sequencing system (Illumina). mRNA-seq samples were pooled and sequenced using Illumina TruSeq Stranded mRNA Library Prep and paired-end sequencing. The samples have 20M pass filter reads with more than 92% of bases above the quality score of Q30. Reads from the samples were trimmed for adapters and low-quality bases using Trimmomatic software before alignment with the reference genome (mm10) and the annotated transcripts using STAR. The mapping statistics were calculated using Picard software. Library complexity was measured in terms of unique fragments in the mapped reads using Picard’s MarkDuplicate utility. The gene expression quantification analysis was performed for all samples using STAR/RSEM tools. Then the differentially expressed genes were analyzed by Limma pipeline with +/-2 fold and p-value of 0.05. Pathway and GO analyses were performed by Gene GO web portal (https://portal.genego.com/cgi/data_manager.cgi). Raw data files are available at the NCBI Gene Expression Omnibus (GEO) server under the accession number GSE148222.

### ChIP-seq

WT and Brd4KO cells (∼ 5 x10^6^ in 15-cm plate) were fixed with 1% formaldehyde (Sigma) in buffer A (50 mM HEPES, pH 7.5, 100 mM NaCl, 1 mM EDTA, and 0.5 mM EGTA) for 10 min and quenched with 0.125 M glycine for 5 min. Cells were washed and incubated in nucleus isolation buffer (50 mM HEPES, pH 7.9, 140 mM NaCl, 1 mM EDTA, 10% Glycerol, 0.5% NP-40, 0.25 % Triton x-100). Nuclear pellets (∼2 x 10^7^) were re-suspended in 1 ml shearing buffer (1 x TE, pH 8, and 0.1% SDS) and transferred to Covaris millitubes and sonicated in Covaris ME200 sonicator. Sonicated chromatin was precipitated with anti-BRD4 antibody (IgG) pre-bound to Dynabeads Protein G (Thermo Fisher) (Dey et al., 2019).

Immunoprecipitated DNA was de-crosslinked, digested with proteinase K, and purified using QIAquick PCR Purification Kit (Qiagen). Precipitated DNA was validated by qChIP. Precipitated DNA (∼ 20 ng) was used for library construction using NEBNext Ultra II DNA Library Prep Kit for Illumina (New England Biolab). Library DNA with fragment length of 200-400 bp was subjected to single end sequencing on a Nextseq system (Illumina). For data analysis, ChIP-seq reads were mapped to the mouse reference genome (mm10) using Bowtie2 v.2.3.4.1 with the following parameters:--sensitive-local. The clonal reads from the aligned reads were removed using PCARD MarkDuplicates and REMOVE_DUPLICATES=true for further analysis. The peak calling was performed using macs2/2.1.1.20160309 program using the following command: macs2 callpeak-t $i.bam-c $i.input.bam -g mm -n $i --outdir $i-Macs-BroadPeak -B -f BAM -- broad -- keep-dup auto --nomodel --extsize 200. The bam files were converted to 1x sequence depth normalized bigWig files as reads per genomic content (RPGC) using deeptools v2.5.46 15 bamCoverage command. For the clustering of cell cycle genes based on BRD4 binding profiles, we used deepTools computeMatrix command to calculate average gene body distribution of BRD4 using the command scale-regions -p 10 -R All-Cell-Cycle2k.bed NonBrd41602.bed Brd4-Cell-564Cycle.bed -S *.bw -b 5000 -a 5000 --regionBodyLength 8000 -- skipZeros -o Brd4-Cycle_scaled.gz--outFileNameMatrix Brd4-Cycle_scaled.tab -- outFileSortedRegions Brd4-Cell_genes.bed. The resulting matrix file was subjected to K-means clustering using plotHeatmap tool and the value of K was chosen at 4. The bed coordinates from the Clusters (C1-C4) were used to predict the genes in each cluster. GO analysis was performed for genes in each cluster using Metascape and GO terms are ranked based on *p*-values. The raw data are submitted under the following accession number GSE153572.

### Acidic extraction of histones

Acidic extraction of histones was performed as reported (https://www.abcam.com/protocols/histone-extraction-protocol-for-western-blot). Briefly, WT and KO MEF cells were harvested at several time points (0, 8, 10, 12, 16, 18, 20 and 24 hrs.) after serum starvation induced synchronization. First, cells were washed with ice cold PBS. Subsequently equal number of cells were lysed in triton extraction buffer (PBS containing 0.5 % Triton X 100 (v/v), 2 mM phenylmethylsulfonyl fluoride (PMSF), 0.02 % (w/v) NaN_3_) for 10 min on ice. Nuclei pellets were collected by centrifugation at 4 ^0^C for 10 min at 6500 g. Nuclei were further washed in triton extraction buffer. Washed nuclei were resuspended in equal amount of 0.2N HCl and acid extracted overnight at 4 ^0^C. Histones (supernatant) were collected by centrifugation at 4 ^0^C for 10 min at 6500 g. Acid extracted histones were further neutralized with 2M NaOH. Histones were separated on NuPAGE^TM^12 % Bis-Tris gel. Two μg of extracts was separated on 12 % SDS-PAGE. Gels were stained with SYPRO Orange (Molecular Probers), photographed and quantified using the ChemiDoc MP Imaging System and its accompanying software, Imaging Lab 5.2 (Bio-Rad).

### Immunoblotting

For detection of other proteins, synchronized WT and BRD4KO cells were harvested at several time points (0, 8, 16, and 24 h.) after serum starvation for synchronization (72h), nuclear extracts was performed as reported (Abmayr et al., 2006). Twenty μg of extracts was separated on 4–20% SDS-PAGE, transferred to polyvinylidene difluoride (PVDF) membranes (Millipore), blocked in 5% of milk in PBS with 0.1% Tween-20 and incubated at 4°Cwith primary antibodies overnight.

Membranes were washed and incubated with horseradish peroxidase-conjugated secondary antibody (1:2000) for 1h, then incubated with ECL (Pierce or GE Healthcare) and imaged using Ecomax X-Ray Film Processor (Protec) or by Azure c600 Biosystems. The Following antibodies with indicated dilution were used by the standard procedures: ATM (ab78, Abcam, 1:1000 dilution), ATR (Santa Cruz Biotechnology Inc, sc-515173, 1:500), Foxm1 (ab180710, Abcam 1: 1000), PLK1 (ab61118, Abcam 1:1000), CDK1(ab133327, Abcam 1:2000), CDK2 (ab32147, Abcam, 1: 2000), CENP F (ab5, Abcam, 1:1000),AUROR A (ab13824, Abcam, 1:1000), AUROR B (PA5_14075, Thermo Fisher Scientific, 1:1000), CENPA (bs-2753R, Bioss.com, 1:500), PARP1 (Cell Signal.com 9542, 1:1000), pATM (Cell Signal.com 5883S 1:1000), pATR (Cell Signal.com 2348S, 1:1000), CHK1 (Cell Signal.com 2360S 1:1000 ab13824, Abcam? 1:1000), ChK2 (Cell Signal.com 2662S 1:1000), H3pS10 (Sigma-Aldrich, 06-507 1:1000), CREST (Protintech #12439-1-A9) and β-ACTIN (8H10D10) (#3700 Cell Signal.com), Topbp1 (Invitrogen #PA576824, 1:1000 dilution), Top1 (Invitrogen #MA5-32228, 1:1000 dilution), Top2a (ThermoFisher. #MA5-38371, 1:500 dilution), RNASEH1 (ThermoFisher. #15606-1-AP, 1:500 dilution), RNASEH2C (abcam ab89726, 1:1000 dilution). Blots were normalized against the loading control (²-ACTIN) and subsequently quantified using Image J software.

### Immunostaining

For immunostaining, cells grown on poly-l-lysine-coated coverslips were fixed in 4% paraformaldehyde for 10 min at room temperature, permeabilized with methanol for 10 min, and stained with primary antibodies for γH2AX (1: 500 #9718 Cell Signal.com),) BRD4 (1:250), H3pS10 (1:500), overnight at 4C . After washing cells were incubated in alexafluor secondary antibodies for 1h and counterstained with Hoechst 33342 (5 ng/ml). For detection of nuclear DNA-RNA hybrid cells were fixed with 100% methanol for 10 min at −20 and were blocked in 2% BSA in PBS overnight at 4C and incubated in primary antibody for mouse S9.6 (1:200) (#ENH001 Kerafast) and rabbit Nucleolin (#22758 Abcam) Sollier et al 2014. Images were captured with Zeiss 880 confocal microscope with 40X water 1.2NA objective. From the captured images intensity per nuclei were determined using ImageJ software (Lam et al., 2020). Statistical significance of fluorescence intensity between experimental condition was determined by two-tailed unpaired t-test using Graph Pad Prizm 9.4.0 software. To determine nuclear S9.6 signal, nuclei were overlaid with Hoechst signal (denoted by white circles in Fig 6) and then intensity were determined by nucleolin subtracted remaining signal of S9.6 using ImageJ.

### Time lapse microscopy and live cell imaging

WT and Brd4KO cells were transduced with a retroviral vector containing H3.1-GFP (Sarai et al., 2013). Cells were incubated in Tokai Hit stage-top incubator at 5% CO2 and ∼100% humidity, with both the chamber and microscope objective (Nikon Plan Apo l 60x NA1.4 oil) warmed to 37°C. Cells were imaged for approximately 18 hours using Yokogawa CSU-W1 spinning disc confocal system (50um pinhole radius) on Nikon Ti2 Eclipse microscope. Sample was illuminated with AOTF-controlled 488nm laser (5% relative power) and fluorescence was detected through FITC emission filter (525/50nm) using Hamamatsu C13440 (Orca Flash 4.0) sCMOS camera. Z-stack (13 x 1um steps, 200msec. exposure/step) was acquired once every 300 sec. for each of the 9 well-separated areas across the cell culture sample, and maximum intensity projection (NIS software) was used to flatten Z-stacks for analysis as well as image and movie preparation (ImageJ).

### Comet assay

Comet assay was performed with Comet Assay Electrophoresis System (Trevigen) in alkaline condition following manufacturer’s recommendation with some modifications (Olive and Banath, 2006; Uehara et al, 2018). Brd4KO, WT or WT Cells treated with ARV-825 were synchronized by serum starvation in serum-free DMEM medium for 24h, and subsequently harvested, washed, and suspended in ice-cold PBS at 2.0 × 10^5^ cells/ml. LMAgarose (Trevigen) at 37°C was added to suspended cells at a ratio of 10:1 (v/v), and the mixture was spread over sample area of comet slides (Trevigen). Slides were kept in dark at 4°C for 10 min, and then immersed in alkaline lysis solution containing 10 mM Tris-HCl (pH 9.5), 2.5 M NaCl, 0.1 M EDTA and 1% (v/v) Triton X-100 at 4°C for 18 hours, and equilibrated in in electrophoresis buffer (0.2 M NaOH and 1 mM EDTA,) for 20 min at 4°C. Electrophoresis was performed at 21 V for 15 min at 4°C. After electrophoresis, slides were washed twice in dH2O for 5 min and 70% ethanol for 5 min then dried at 37°C for 30 min. The DNA was stained with SYBR-Gold (1ul in 10 ml TE buffer) for 30 min at room temperature. Slides were briefly rinsed with dH2O and dried at 37°C. Comet images were captured with Zeiss 880 confocal microscope with 40X water 1.2NA objective. Tail moments were determined using Image J software with open comet plugin (Gyori et al, 2014; Lam et al., 2020).

## Supporting information

Tiyun Wu et al. Supplemental information

Tiyun Wu et al. Supplemental figures

Tiyun Wu et al. Table S1-2023

Tiyun Wu et al. Table S2-2023

Tiyun Wu et al. Table S3-2023

Tiyun Wu et al. Table S4-2023

Tiyun Wu et al. Table S5-2023

## ACKNOWLEDGEMENTS

The authors thank J. Cooper, D. Singer, C. Wu (NCI) for advice, discussions, and critical reading of the manuscript. We thank Dr. Crouch and Uehara for providing instrument and information for comet assays. We also thank Paula Maia for technical assistance, members of the Ozato lab for technical advice. This work was supported by the NICHD Intramural programs ZIA HD008015-13.

## AUTHOR CONSTRIBUTIONS

KO conceived the study, steered the project along with TW. TW performed FACS, RNA-seq, ChIPseq, RT-qPCR, immunoblot and immunostaining for H2AX and S9.6, comet assay. HH carried out S and G2/M block study. XC analyzed RNA-seq data. MB and RP analyzed ChIP-seq data. JW conducted time lapse imaging and analyses. CC performed apoptosis assay. AD performed ARV-825 treated cells, immunostaining for H3pS10, pATR, ATM and comet assay. FK worked in WI38 cell and performed parelle work as in MEFs. SC performed and analysis with core histone proteins and immunoblot for H3pS10. XH analysis with core histone proteins and gave general guidance throughout the work. TW and KO prepared figures and wrote the manuscript.

## COMPETING FINANCIAL INTERSTS

All authors declare no competing financial interests.

## REFERENCES

1. Barrows, J. K. Lin, B. Quaas, C. E. Fullbright, G. Wallace, E. N. Long, D. T. (2022). BRD4 promotes resection and homology-directed repair of DNA double-strand breaks. Nat Commun 13, 3016

2. Behera, V. Stonestrom, A. J. Hamagami, N. Hsiung, C. C. Keller, C. A. Giardine, B. Sidoli, S. Yuan, Z. F. Bhanu, N. V. Werner, M. T. Wang, H. Garcia, B. A. Hardison, R. C. Blobel, G. A. (2019). Interrogating Histone Acetylation and BRD4 as Mitotic Bookmarks of Transcription. Cell Rep 27, 400–415 e5

3. Bertoli, C., Skotheim, J.M., and de Bruin, R.A. (2013). Control of cell cycle transcription during G1 and S phases. Nat Rev Mol Cell Biol 14, 518–528.

4. Blackford, A.N., and Jackson, S.P. (2017). ATM, ATR, and DNA-PK: The Trinity at the Heart of the DNA Damage Response. Mol Cell 66, 801–817.

5. Bonner, W.M., Redon, C.E., Dickey, J.S., Nakamura, A.J., Sedelnikova, O.A., Solier, S., and Pommier, Y. (2008). GammaH2AX and cancer. Nat Rev Cancer 8, 957–967.

6. Bou-Nader, C. Bothra, A. Garboczi, D. N. Leppla, S. H. Zhang, J. (2022). Structural basis of R-loop recognition by the S9.6 monoclonal antibody. Nat Commun 13, 1641

7. Casella, G. Munk, R. Kim, K. M. Piao, Y. De, S. Abdelmohsen, K. Gorospe, M. (2019). Transcriptome signature of cellular senescence. Nucleic Acids Res 47, 11476

8. Choi, E. H. Yoon, S. Park, K. S. Kim, K. P. (2017). The Homologous Recombination Machinery Orchestrates Post-replication DNA Repair During Self-renewal of Mouse Embryonic Stem Cells. Sci Rep 7, 11610

9. Costa, R.H. (2005). FoxM1 dances with mitosis. NATURE CELL BIOLOGY 7, 108–110.

10. Crasta K, Ganem NJ, Dagher R, Lantermann AB, Ivanova EV, Pan Y, Nezi L, Protopopov A, Chowdhury D, Pellman D. (2012). DNA breaks and chromosome pulverization from errors in mitosis. Nature. 482, 53-58.

11. Delmore, J.E., Issa, G.C., Lemieux, M.E., Rahl, P.B., Shi, J., Jacobs, H.M., Kastritis, E., Gilpatrick, T., Paranal, R.M., Qi, J., et al. (2011). BET bromodomain inhibition as a therapeutic strategy to target c-Myc. Cell 146, 904–917.

12. Devaiah BN, Singer DS. (2013). Two faces of brd4: mitotic bookmark and transcriptional lynchpin. Transcription. 4, 13–17.

13. Devaiah, B.N., Case-Borden, C., Gegonne, A., Hsu, C.H., Chen, Q., Meerzaman, D., Dey, A., Ozato, K., and Singer, D.S. (2016). BRD4 is a histone acetyltransferase that evicts nucleosomes from chromatin. Nat Struct Mol Biol 23, 540–548.

14. Dey, A., Chitsaz, F., Abbasi, A., Misteli, T. & Ozato, K. (2003). The double bromodomain protein Brd4 binds to acetylated chromatin during interphase and mitosis. Proc Natl Acad Sci U S A 100, 8758–8763.

15. Dey, A., Ellenberg, J., Farina, A., Coleman, A.E., Maruyama, T., Sciortino, S., Lippincott-Schwartz, J., and Ozato, K. (2000). A bromodomain protein, MCAP, associates with mitotic chromosomes and affects G(2)-to-M transition. Mol Cell Biol 20, 6537-6549.

16. Dey, A., Nishiyama, A., Karpova, T., McNally, J., and Ozato, K. (2009). Brd4 marks select genes on mitotic chromatin and directs postmitotic transcription. Mol Biol Cell 20, 4899–4909.

17. Dey, A., Yang, W., Gegonne, A., Nishiyama, A., Pan, R., Yagi, R., Grinberg, A., Finkelman, F.D., Pfeifer, K., Zhu, J., et al. (2019). BRD4 directs hematopoietic stem cell development and modulates macrophage inflammatory responses. EMBO J 38.

18. Donati, B., Lorenzini, E. & Ciarrocchi, A. (2018). BRD4 and Cancer: going beyond transcriptional regulation. Mol Cancer. 17, 164–177.

19. Edwards, D. S. Maganti, R. Tanksley, J. P. Luo, J. Park, J. J. H. Balkanska-Sinclair, E. Ling, J. Floyd, S. R. (2020) BRD4 Prevents R-Loop Formation and Transcription-Replication Conflicts by Ensuring Efficient Transcription Elongation. Cell Rep 32, 108–166.

20. Engeland, K. (2022) Cell cycle regulation: p53-p21-RB signaling. Cell Death Differ 29, 946–960

21. Filippakopoulos, P., Qi, J., Picaud, S., Shen, Y., Smith, W.B., Fedorov, O., Morse, E.M., Keates, T., Hickman, T.T., Felletar, I., et al. (2010). Selective inhibition of BET bromodomains. Nature 468, 1067-1073.

22. Floyd, S.R., Pacold, M.E., Huang, Q., Clarke, S.M., Lam, F.C., Cannell, I.G., Bryson, B.D., Rameseder, J., Lee, M.J., Blake, E.J., et al. (2013). The bromodomain protein Brd4 insulates chromatin from DNA damage signalling. Nature 498, 246–250.

23. >Galgano, P. J. Schildkraut, C. L. (2006). G1/S Phase Synchronization using Double Thymidine Synchronization. CSH Protoc 2006 Jul 1;2006(2):pdb.prot4487.

24. Grant, G.D., Brooks, L3rd., Zhang, X., Mahoney, J.M., Martyanov, V., Wood, T.A., Sherlock, G., Cheng, C., and Whitfield, M.L. (2013). Identification of cell cycle-regulated genes periodically expressed in U2OS cells and their regulation by FOXM1 and E2F transcription factors. Mol Biol Cell 24, 3634-3650.

25. Gyori, B. M. Venkatachalam, G. Thiagarajan, P. S. Hsu, D. Clement, M. V. (2014). OpenComet: an automated tool for comet assay image analysis. Redox Biol 2, 457–465.

26. Hargreaves, D.C., Horng, T., and Medzhitov, R. (2009). Control of inducible gene expression by signal-dependent transcriptional elongation. Cell 138, 129–145.

27. Hayflick, L. (1965). The Limited in Vitro Lifetime of Human Diploid Cell Strains. Exp Cell Res 37, 614–636.

28. Hegazy, Y. A. Fernando, C. M. Tran, E. J. (2020). The balancing act of R-loop biology: The good, the bad, and the ugly. J Biol Chem 295, 905–913.

29. Hnisz, D., Abraham, B.J., Lee, T.I., Lau, A., Saint-Andre, V., Sigova, A.A., Hoke, H.A., and Young, R.A. (2013). Super-enhancers in the control of cell identity and disease. Cell 155, 934–947

30. Houzelstein D, Bullock SL, Lynch DE, Grigorieva EF, Wilson VA, Beddington RS. (2002). Growth and Early Postimplantation Defects in Mice Deficient for theBromodomain-Containing Protein Brd4. Molecular and Cellular Biology 22, 3794–3802.

31. Jang, M.K., Mochizuki, K., Zhou, M., Jeong, H.S., Brady, J.N., and Ozato, K. (2005). The bromodomain protein Brd4 is a positive regulatory component of P-TEFb and stimulates RNA polymerase II-dependent transcription. Mol Cell 19, 523–534.

32. Kabeche, L., Nguyen, H.D., Buisson, R., and Zou, L. (2018). A mitosis-specific and R loop-driven ATR pathway promotes faithful chromosome segregation. Science 359, 108–114.

33. Kanno, T., Kanno, Y., LeRoy, G., Campos, E., Sun, H.W., Brooks, S.R., Vahedi, G., Heightman, T.D., Garcia, B.A., Reinberg, D., et al. (2014). BRD4 assists elongation of both coding and enhancer RNAs by interacting with acetylated histones. Nat Struct Mol Biol 21, 1047–1057.

34. Kim, J. J. Lee, S. Y. Gong, F. Battenhouse, A. M. Boutz, D. R. Bashyal, A. Refvik, S. T. Chiang, C. M. Xhemalce, B. Paull, T. T. Brodbelt, J. S. Marcotte, E. M. Miller, K. M. (2019). Systematic bromodomain protein screens identify homologous recombination and R-loop suppression pathways involved in genome integrity. Genes Dev 33, 1751–1774.

35. Lam, F. C. Kong, Y. W. Huang, Q. Vu Han, T. L. Maffa, A. D. Kasper, E. M. Yaffe, M. B. (2020). BRD4 prevents the accumulation of R-loops and protects against transcription-replication collision events and DNA damage. Nat Commun 11, 4083–4102.

36. Laoukili, J., Kooistra, M.R., Bras, A., Kauw, J., Kerkhoven, R.M., Morrison, A., Clevers, H., and Medema, R.H. (2005). FoxM1 is required for execution of the mitotic programme and chromosome stability. Nat Cell Biol 7, 126–136.

37. Lewis, C.W., and Golsteyn, R.M. (2016). Cancer cells that survive checkpoint adaptation contain micronuclei that harbor damaged DNA. Cell Cycle 15, 3131–3145.

38. Li, X., Baek, G., Ramanand, S.G., Sharp, A., Gao, Y., Yuan, W., Welti, J., Rodrigues, D.N., Dolling, D., Figueiredo, I., et al. (2018). BRD4 Promotes DNA Repair and Mediates the Formation of TMPRSS2-ERG Gene Rearrangements in Prostate Cancer. Cell Rep 22, 796–808.

39. Liu, Y. Nielsen, C. F. Yao, Q. Hickson, I. D. (2014) The origins and processing of ultra fine anaphase DNA bridges. Curr Opin Genet Dev 26, 1–5.

40. Loven, J., Hoke, H.A., Lin, C.Y., Lau, A., Orlando, D.A., Vakoc, C.R., Bradner, J.E., Lee, T.I., and Young, R.A. (2013). Selective inhibition of tumor oncogenes by disruption of super-enhancers. Cell 153, 320–334.

41. Marzluff, W.F., Gongidi, P., Woods, K.R., Jin, J., and Maltais, L.J. (2002). The human and mouse replication-dependent histone genes. Genomics 80, 487–498.

42. Mashimo, M. Onishi, M. Uno, A. Tanimichi, A. Nobeyama, A. Mori, M. Yamada, S. Negi, S. Bu, X. Kato, J. Moss, J. Sanada, N. Kizu, R. Fujii, T. (2021). The 89-kDa PARP1 cleavage fragment serves as a cytoplasmic PAR carrier to induce AIF-mediated apoptosis. J Biol Chem 296, 100046

43. Mochizuki, K., Nishiyama, A., Jang, M.K., Dey, A., Ghosh, A., Tamura, T., Natsume, H., Yao, H., and Ozato, K. (2008). The bromodomain protein Brd4 stimulates G1 gene transcription and promotes progression to S phase. J Biol Chem 283, 9040-9048.

44. Nicodeme, E., Jeffrey, K.L., Schaefer, U., Beinke, S., Dewell, S., Chung, C.W., Chandwani, R., Marazzi, I., Wilson, P., Coste, H., et al. (2010). Suppression of inflammation by a synthetic histone mimic. Nature 468, 1119–1123.

45. Niehrs, C. Luke, B. (2020). Regulatory R-loops as facilitators of gene expression and genome stability. Nat Rev Mol Cell Biol 21, 167–178.

46. Nishiyama, A., Dey, A., Miyazaki, J. Ozato K. (2006). Brd4 Is Required for Recovery from Antimicrotubule Drug-induced Mitotic Arrest: Preservation of Acetylated Chromatin. Molecular Cell Biology 17, 814–823.

47. Nishiyama, A., Dey, A., Tamura, T., Ko, M., and Ozato, K. (2012). Activation of JNK triggers release of Brd4 from mitotic chromosomes and mediates protection from drug-induced mitotic stress. PLoS One 7, e34719.

48. Nizami, Z., Deryusheva, S., and Gall, J.G. (2010). The Cajal body and histone locus body. Cold Spring Harb Perspect Biol 2, a000653.

49. Olive, P. L. Banath, J. P. (2006). The comet assay: a method to measure DNA damage in individual cells. Nat Protoc 1, 23–29.

50. Palazzo, L., Della Monica, R., Visconti, R., Costanzo, V., and Grieco, D. (2014). ATM controls proper mitotic spindle structure. Cell Cycle 13, 1091–1100.

51. Patel, M. C. Debrosse, M. Smith, M. Dey, A. Huynh, W. Sarai, N. Heightman, T. D. Tamura, T. Ozato,K. (2013). BRD4 coordinates recruitment of pause release factor P-TEFb and the pausing complex NELF/DSIF to regulate transcription elongation of interferon-stimulated genes. Mol Cell Biol 33, 2497–507.

52. Pelish, H.E., Liau, B.B., Nitulescu, II, Tangpeerachaikul, A., Poss, Z.C., Da Silva, D.H., Caruso, B.T., Arefolov, A., Fadeyi, O., Christie, A.L., et al. (2015). Mediator kinase inhibition further activates super-enhancer-associated genes in AML. Nature 526, 273–276.

53. Promonet, A. Padioleau, I. Liu, Y. Sanz, L. Biernacka, A. Schmitz, A. L. Skrzypczak, M. Sarrazin, A. Mettling, C. Rowicka, M. Ginalski, K. Chedin, F. Chen, C.L. Lin, Y. L. Pasero, P. (2020). Topoisomerase 1 prevents replication stress at R-loop-enriched transcription termination sites. Nat Commun 11, 3940

54. Raina, K. Lu, J. Qian, Y. Altieri, M. Gordon, D. Rossi, A. M. Wang, J. Chen, X. Dong, H. Siu, K. Winkler, Crew,A. P. Crews, C. M Coleman, K. G. (2016). PROTAC-induced BET protein degradation as a therapy for castration-resistant prostate cancer. Proc Natl Acad Sci U S A 1 13, 7124–7129.

55. Ray Chaudhuri, A. Nussenzweig, A. (2017). The multifaceted roles of PARP1 in DNA repair and chromatin remodelling. Nat Rev Mol Cell Biol 18, 610–621.

56. Saldivar, J., Hamperl, S., Bocek, M.J., Chung, M., Bass, T.E., Cisneros-Soberanis, F., Samejima, K., Xie, L., Paulson, J.R., Earnshaw, W.C., et al. (2018). An intrinsic S/G2 checkpoint enforced by ATR. Science 361, 806–810.

57. Santaguida, S., and Amon, A. (2015). Short-and long-term effects of chromosome mis-segregation and aneuploidy. Nat Rev Mol Cell Biol 16, 473–485.

58. Sarai, N., Nimura, K., Tamura, T., Kanno, T., Patel, M.C., Heightman, T.D., Ura, K., and Ozato, K. (2013). WHSC1 links transcription elongation to HIRA-mediated histone H3.3 deposition. EMBO J 32, 2392–2406.

59. Shechter, D., Dormann, H.L., Allis, C.D., and Hake, S.B. (2007). Extraction, purification and analysis of histones. Nat Protoc 2, 1445-1457.

60. Shi, J., and Vakoc, C.R. (2014). The mechanisms behind the therapeutic activity of BET bromodomain inhibition. Mol Cell 54, 728–736.

61. Skoufias, D.A., DeBonis, S., Saoudi, Y., Lebeau, L., Crevel, I., Cross, R., Wade, R.H., Hackney, D., and Kozielski, F. (2006). S-trityl-L-cysteine is a reversible, tight binding inhibitor of the human kinesin Eg5 that specifically blocks mitotic progression. J Biol Chem 281, 17559–17569.

62. Sollier, J., and Cimprich, K. A. (2015). Breaking bad: R-loops and genome integrity. Trends Cell Biol 25, 514–22.

63. Sollier, J., Stork, C. T. Garcia-Rubio, M. L. Paulsen, R. D. Aguilera, A. Cimprich, K. A. (2014). Transcription-coupled nucleotide excision repair factors promote R-loop-induced genome instability. Mol Cell 56, 777–85.

64. Sollier, J. Cimprich, K. A. (2015). Breaking bad: R-loops and genome integrity. Trends Cell Biol 25, 514–22.

65. Uehara, R. Cerritelli, S. M. Hasin, N. Sakhuja, K. London, M. Iranzo, J. Chon, H. Grinberg, A. Crouch, R. J. (2018). Two RNase H2 Mutants with Differential rNMP Processing Activity Reveal a Threshold of Ribonucleotide Tolerance for Embryonic Development. Cell Rep 25, 1135–1145 e5.

66. Walczak, C. E. Cai, S. Khodjakov, A. (2010). Mechanisms of chromosome behaviour during mitosis. Nat Rev Mol Cell Biol 11, 91–102.

67. Wei, Y. Yu, L. Bowen, J. Gorovsky, M. A. Allis, C. D. (1999). Phosphorylation of histone H3 is required for proper chromosome condensation and segregation. Cell 97, 99–109.

68. White, M.E., Fenger, J.M., and Carson, W.E3rd., (2019). Emerging roles of and therapeutic strategies targeting BRD4 in cancer. Cell Immunol 337, 48-53.

69. Whyte, W.A., Orlando, D.A., Hnisz, D., Abraham, B.J., Lin, C.Y., Kagey, M.H., Rahl, P.B., Lee, T.I., and Young, R.A. (2013). Master transcription factors and mediator establish super-enhancers at key cell identity genes. Cell 153, 307–319.

70. Wierstra, I. (2013). The transcription factor FOXM1 (Forkhead box M1): proliferation-specific expression, transcription factor function, target genes, mouse models, and normal biological roles. Adv Cancer Res 118, 97–398.

71. Williams, A. B. Schumacher, B. (2016). P53 in the DNA-Damage-Repair Process. Cold Spring Harb Perspect Med 6, 1–15.

72. Wu, S.Y., and Chiang, C.M. (2007). The double bromodomain-containing chromatin adaptor Brd4 and transcriptional regulation. J Biol Chem 282, 13141–13145.

73. Ye, X., Wei, Y., Nalepa, G., and Harper, J.W. (2003). The cyclin E/Cdk2 substrate p220(NPAT) is required for S-phase entry, histone gene expression, and Cajal body maintenance in human somatic cells. Mol Cell Biol 23, 8586–8600.

74. Zhang, J., Dulak, A.M., Hattersley, M.M., Willis, B.S., Nikkila, J., Wang, A., Lau, A., Reimer, C., Zinda, M., Fawell, S.E., et al. (2018). BRD4 facilitates replication stress-induced DNA damage response. Oncogene 37, 3763–3777.

75. Zhang, W., Prakash, C., Sum, C., Gong, Y., Li, Y., Kwok, J.J., Thiessen, N., Pettersson, S., Jones, S.J., Knapp, S., et al. (2012). Bromodomain-containing protein 4 (BRD4) regulates RNA polymerase II serine 2 phosphorylation in human CD4+ T cells. J Biol Chem 287, 43137–43155.

76. Zuber, J., Shi, J., Wang, E., Rappaport, A.R., Herrmann, H., Sison, E.A., Magoon, D., Qi, J., Blatt, K., Wunderlich, M., et al. (2011). RNAi screen identifies Brd4 as a therapeutic target in acute myeloid leukaemia. Nature 478, 524–528.

