## Supplementary material for "BRD4 directs mitotic cell division by inhibiting DNA damage": Tiyun Wu et al. Supplemental information

**Supplementary information**

Reporting Summary

**Supplemental Figure S1 to S6**

Supplemental Figure S1, related to Figure 1.

Brd4 knockout affects cell cycle progression

Supplemental Figure S2, related to Figure 2.

RNA-seq analysis of BRD4-dependent cell cycle genes

Supplemental Figure S3, related to Figure 3.
qRT-PCR and quantification of immunoblot in selected genes in Brd4KO cells

Supplemental Figure S4, related to Figure 4.
BRD4 ChIP-seq profiles

Supplemental Figure S5, related to Figure 5.
Effects of drugs that arrest mitotic progression and Movies of mitotic progression of live cells

Supplemental Figure S6, related to Figure 6.
Brd4KO cells incur DNA damage and accumulate R-loops, a model of Brd4 function in cell cycle control

**Supplementary Videos**

Mitotic progression 3 movies (WT, Brd4KO-1 and Brd4KO-2)

**Supplementary Tables 1 to 5**

Table S1

2860 cell cycle genes were identified by comparing expression at 0h to those at later time points throughout the cell cycle (0h/4h, or /8h, 12h, 16h, 20h, 24h) in synchronized WT cells, after removing duplications. Specifically, n=1808 0h-4h, n=1764 0h-8h, n=1422 0h-12h, n=1201 0h-12h, n=995 0h-12h and n=627 0h-24h. The differentially expressed cell cycle genes were analyzed by Limma pipeline with +/-2 fold and p-value of 0.05.

Table S2

2166 Brd4 regulated genes were differentially expressed between WT and Brd4KO cells at each time point (0h, 4h, 8h, 12h, 16h, 20h and 24h). Specifically, n=305 0h, n=321 4h, n=381 8h, n=173 12h, n=424 16h, n=258 20h and n=352 24h. The differentially expressed genes were analyzed by Limma pipeline with +/-2 fold and p-value of 0.05.

Table S3

455 genes were down-regulated in Brd4KO cells.

Table S4

Clusters (C1-C4) were used to predict the genes in each cluster. GO analysis was performed for genes in each cluster using Metascape and GO terms are ranked based on *p*-values

Table S5

Primer sequences used for qPCR
