## Supplementary material for "BRD4 directs mitotic cell division by inhibiting DNA damage": Tiyun Wu et al. Supplemental figures

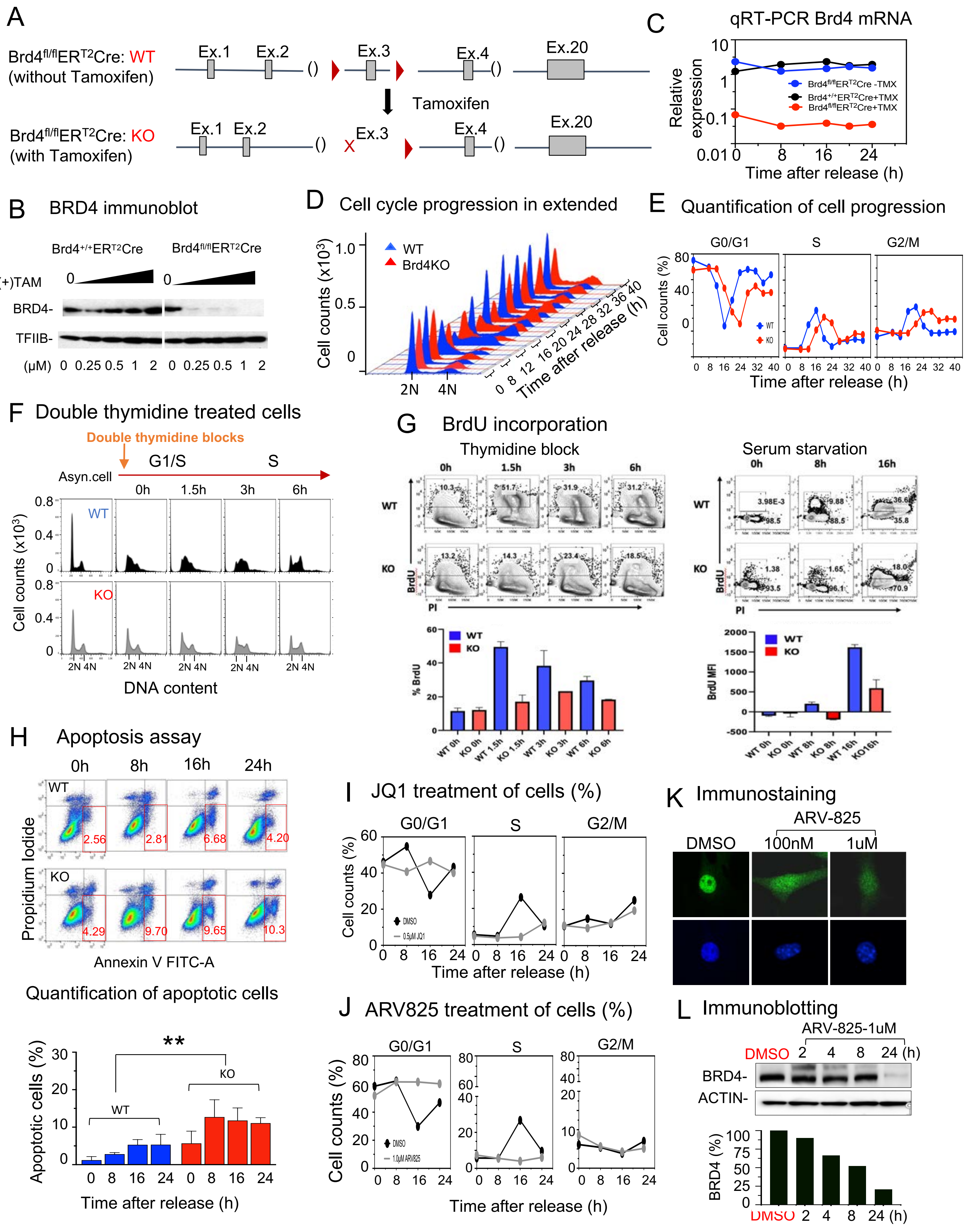
A
Brd4fl/flERT2Cre: WT (without Tamoxifen)
Brd4fl/flERT2Cre: KO (with Tamoxifen)
Ex.1 Ex.2 Ex.3 Ex.4 Ex.20
Tamoxifen
C
qRT-PCR Brd4 mRNA
Relative expression
Time after release (h)
Brd4fl/flERT2Cre -TMX
Brd4+/+ERT2Cre+TMX
Brd4fl/flERT2Cre+TMX
B
BRD4 immunoblot
Brd4+/+ERT2Cre
Brd4fl/flERT2Cre
(+)TAM
BRD4-
TFIIB-
(μM) 0 0.25 0.5 1 2 0 0.25 0.5 1 2
D
Cell cycle progression in extended
WT
Brd4KO
Cell counts (x10³)
Time after release (h)
E
Quantification of cell progression
G0/G1
S
G2/M
Cell counts (%)
Time after release (h)
F
Double thymidine treated cells
Double thymidine blocks
Asyn.cell
G1/S
S
0h 1.5h 3h 6h
WT
KO
Cell counts (x10³)
DNA content
G
BrdU incorporation
Thymidine block
Serum starvation
0h 1.5h 3h 6h
0h 8h 16h
BrdU
PI
% BrdU
BrdU MFI
H
Apoptosis assay
0h 8h 16h 24h
Propidium Iodide
Annexin V FITC-A
Quantification of apoptotic cells
Apoptotic cells (%)
Time after release (h)
I
JQ1 treatment of cells (%)
DMSO
0.5μM JQ1
Time after release (h)
J
ARV825 treatment of cells (%)
DMSO
1.0μM ARV825
Time after release (h)
K
Immunostaining
DMSO
ARV-825
100nM
1uM
L
Immunoblotting
ARV-825-1uM
DMSO 2 4 8 24 (h)
BRD4-
ACTIN-
BRD4 (%)

**Supplementary information, Figure S1**

**Brd4 knockout affects cell cycle progression**

A. Diagram for Brd4 deletion.

B. Loss of BRD4 protein. Brd4$^{+/+}$ ER$^{T2}$Cre cells (left) or Brd4$^{f/f}$ER$^{T2}$Cre cells (right) were treated with Tamoxifen at indicated concentrations for 96h. Twenty µg of nuclear extracts were immunoblotted with rabbit anti-BRD4 antibody. TFIIB was used as a loading control.

C. Brd4$^{f/f}$ER$^{T2}$Cre or Brd4$^{+/+}$ER$^{T2}$Cre cells were treated with (+) or without (-)Tamoxifen, and then synchronized by serum starvation. The cells collected at indicated time after release, Brd4 mRNA were measured by qRT-PCR and normalized by Hprt.

D. WT and Brd4KO cells were synchronized as in Figuer 1C, allowed to proceed for up to 40h and DNA contents were determined as in Figure 1C.

E. Quantification of cells progression at each stage during cell cycle in Figure S1D.

F. WT and Brd4KO cells were treated twice with thymidine (200 µg/ml), released and allowed to proceed up to 6h. Cells were stained with propidium iodide (25 µg/ml) and DNA contents were determined by flow cytometry.

G. WT and Brd4KO cells were treated with thymidine (left) or serum starvation (right) as in Figure S1F or Figure 1C, released and allowed to proceed at indicated time. BrdU (1mM) was added 2h before harvest, cells were stained with propidium iodide (25 µg/ml) and DNA contents were determined by flow cytometry (top). Quantification of gate was set on BrdU positive population (bottom).

H. WT or Brd4KO cells were synchronized as above, stained with Alexa Fluor® 488 Annexin FITC (X axis) and propidium iodide (Y axis) and analyzed by flowcytometry. The percentages of apoptotic cells are shown in the bottom right quadrant (red). Quantification of gate was set apoptotic cells at indicated times and are shown (bottom). Significance was calculated using unpaired t test ($P<0.003$).

I. WT cells synchronized as above were treated with JQ1 as in Figure 1E. The percentages of cells at G0/G1, S or G2/M were estimated as above.

J. WT cells synchronized as above were treated with ARV-825 as in Figure 1E. The percentages of cells at G0/G1, S or G2/M were estimated as above.

K. Immunestaining of BRD4 for ARV-825 treated or DMSO (at indicated concentration) WT cells for 24h.

L. Immunoblot of BRD4 after ARV-825(1µM) treated WT cells at indicated time points (0, 4, 8 and 24h) and control with DMSO (Top). βACTIN was used as a loading control.

**Supplementary information, Figure S2**

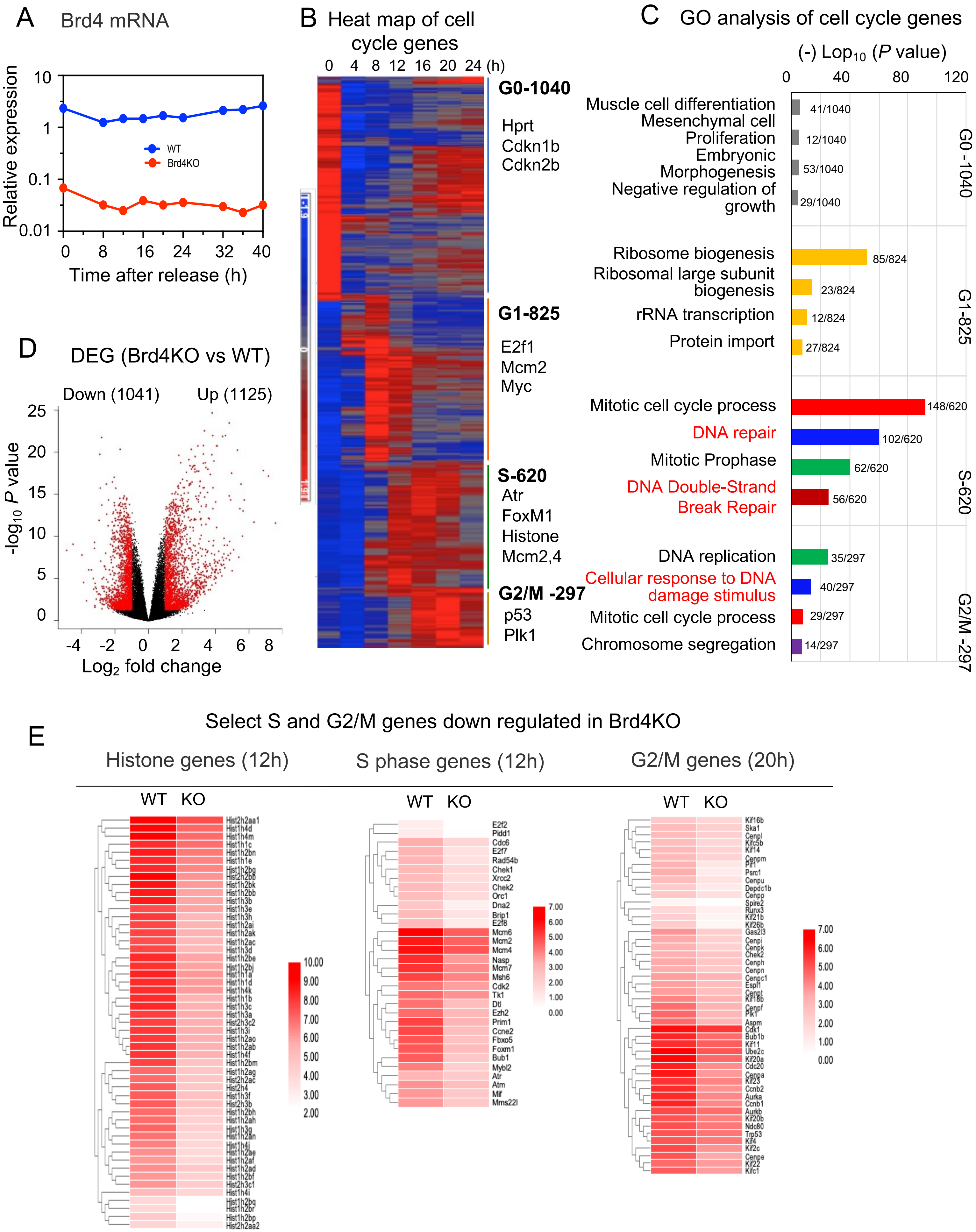

**Supplementary information, Figure S2**

**RNA-seq analysis of BRD4-dependent cell cycle genes**

A. Confirmation of Brd4 loss. WT and Brd4KO cells were synchronized with extended time to release up to 40h as in Fig S1D, and expression of Brd4 mRNA was detected at indicated times by qRT-PCR.

B. Heat map of cells cycle genes identified in WT cells. Included are 1040 G0 genes at 0h, 825 G1 genes peaking at 8h-12h, 620 S genes peaking at 12h-16h and 297 G2/M genes detected at 16h-24h.

C. GO analysis of cell cycles genes identified Fig.S2B. G0, G1, S, G2/M genes were analyzed by Enriched Ontology Cluster program of Metascape.

D. Volcano plots depicting differentially expressed genes between WT and Brd4KO cells. Genes (FC>2, $p$-value < 0.05) at any time point are shown in red. Up and Down indicate the number of genes with higher or lower expression in Brd4KO cells.

E. Heat maps of select S and G2/M genes downregulated in Brd4KO cells. Left. Core histones and histone H1 genes in WT or Brd4KO cells expressed at 12h (S). Expression levels are graded on the scale bar. Middle. Other S phase genes (12h), including Foxm1, Atm, Atr and those involved in DNA replication e.g., Orc1, Mcms (2,6,7). Right. G2/M genes, expressed at 20h, involved in transition from G2 to M e.g., Ccnb1, Cdk1, Cdk2, Cdc20, Bub1, Aurka, Aurkb, Cenps (as Cenpa or f,I,h,n) and Kifs (as Kif1 or 4,6).

### A RT-qPCR

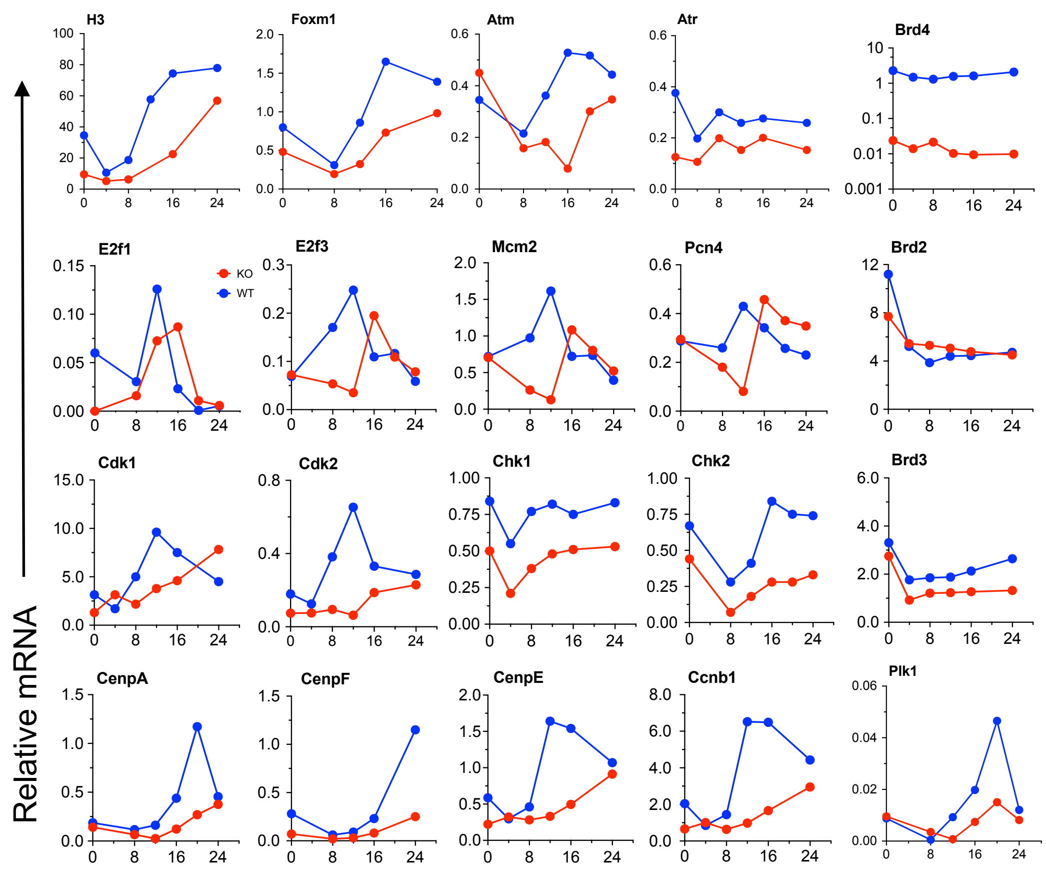


### B Quantification of immunoblots

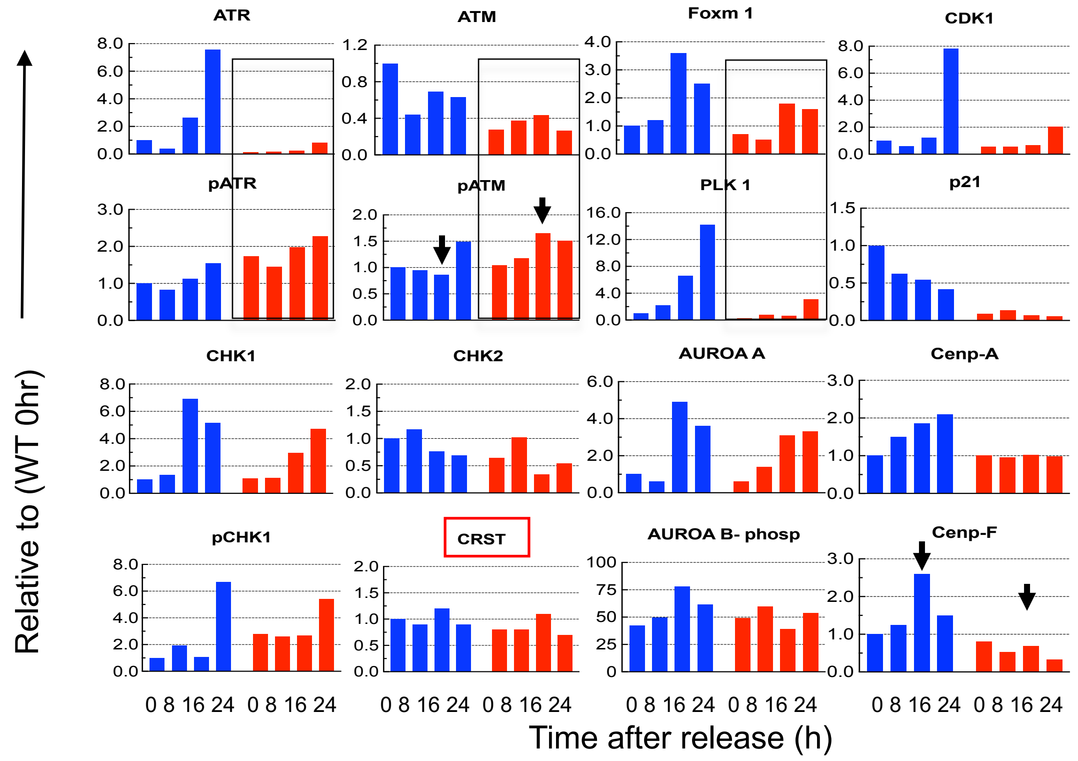

**Supplementary information, Figure S3**

**qRT-PCR and quantification of immunoblot in selected genes in Brd4KO cells**

A. qRT-PCR confirmation of cell cycle genes downregulated in Brd4KO cells. qRT-PCR was performed for indicated cell cycle genes in synchronized WT and Brd4KO cells. mRNA expression was normalized to Hprt. Vales represents the average of three independent experiments.

B. Quantification of Immunoblot data in Fig. 3b. Bands in the blots were normalized against the loading control (β–ACTIN) and quantified using Image J software. CRST, which is not regulated by BRD4, showed very similar patterns of expression in WT vs BRD4KO during cell cycle. Results were obtained from at least two independent experiments.

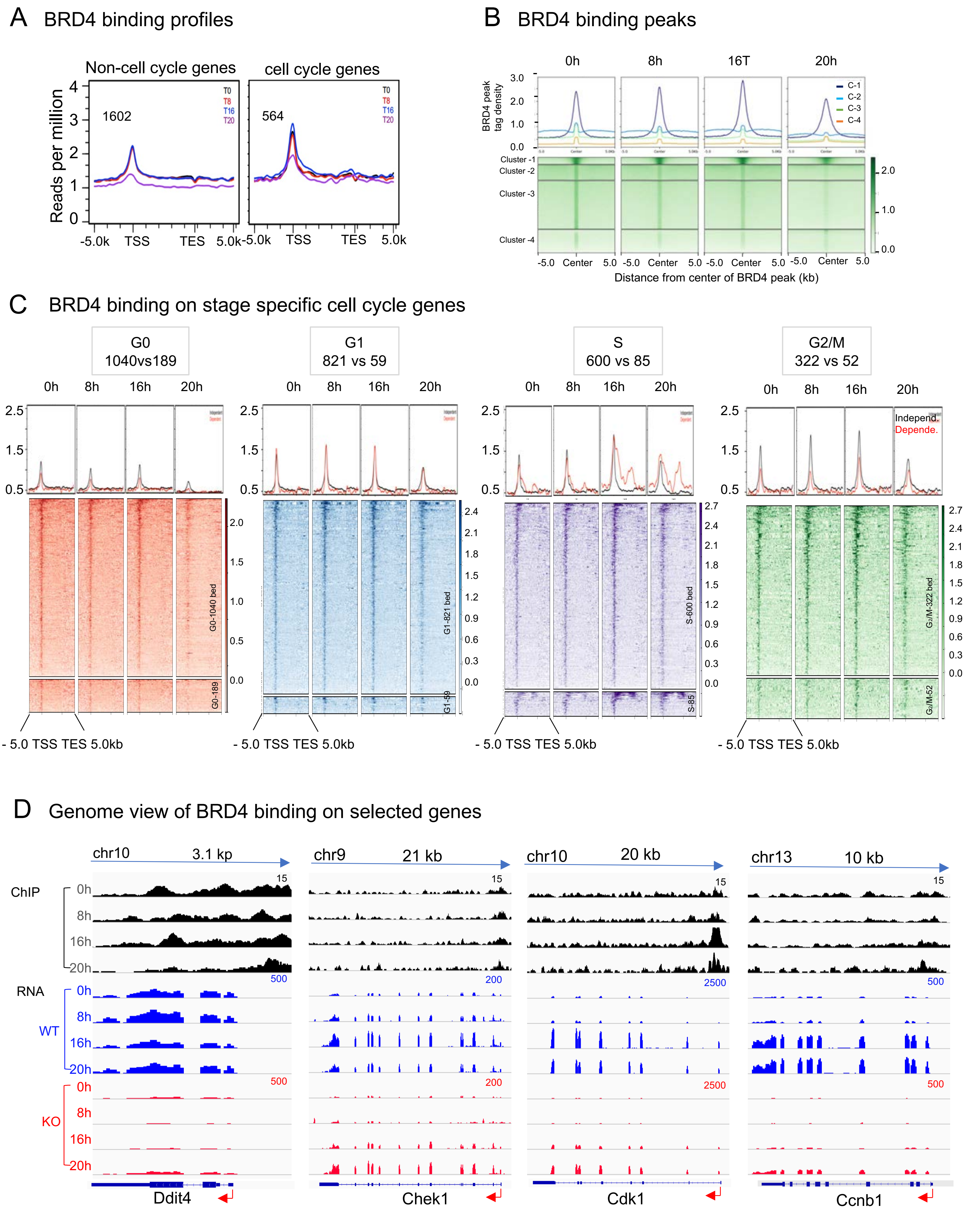
A BRD4 binding profiles
Non-cell cycle genes
cell cycle genes
Reads per million
1602
564
T0
T8
T16
T20
-5.0k TSS TES 5.0k
B BRD4 binding peaks
0h 8h 16T 20h
BRD4 peak tag density
C-1
C-2
C-3
C-4
Cluster -1
Cluster -2
Cluster -3
Cluster -4
-5.0 Center 5.0
Distance from center of BRD4 peak (kb)
C BRD4 binding on stage specific cell cycle genes
G0 1040vs189
G1 821 vs 59
S 600 vs 85
G2/M 322 vs 52
0h 8h 16h 20h
Independ.
Depende.
G0-1040 bed
G0-189
G1-821 bed
G1-59
S-600 bed
S-85
G2/M-322 bed
G2/M-52
- 5.0 TSS TES 5.0kb
D Genome view of BRD4 binding on selected genes
chr10 3.1 kp
chr9 21 kb
chr10 20 kb
chr13 10 kb
ChIP
RNA
WT
KO
0h
8h
16h
20h
Ddit4
Chek1
Cdk1
Ccnb1

**Supplementary information, Figure S4**

**BRD4 ChIP-seq profiles**

A. Average BRD4 distribution is plotted for non-cell cycle genes and cell cycle genes regulated by BRD4.

B. BRD4 signals on cell cycle genes in Fig. 4b was replotted to show the highest BRD4 binding at the center +/- 5Kb distance. Top, the average profiles; bottom, heat maps at indicated times.

C. BRD4 binding on stage specific cell cycle genes. (Top) Average BRD4 binding plotted for all cycle regulated genes, specific for each stage (Figure 2b,e, Figure S2b) at indicated times. Black lines indicate total cell cycle genes, red lines denote BRD4 dependent genes (gene numbers indicated at box on the top). Note constitutive BRD4 occupancy in contrast to stage specific RNA expression. (Bottom) Heat map; Top panels represent BRD4 binding on total cell cycle genes and the bottom panels show BRD4 dependent cell cycle genes, aligned according to BRD4 signal intensity.

D. IGV profiles of BRD4 occupancy (top) and RNA-seq peaks (middle and bottom) for indicated cell cycle gens in WT and Brd4KO cells. Gene names and the exon-intron organization are shown below.

A Drug treatment

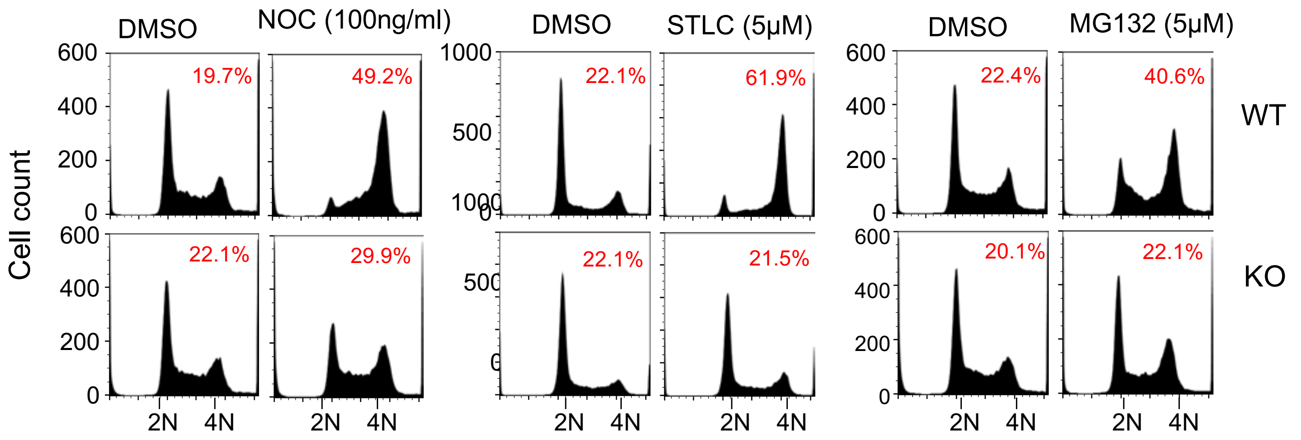


B

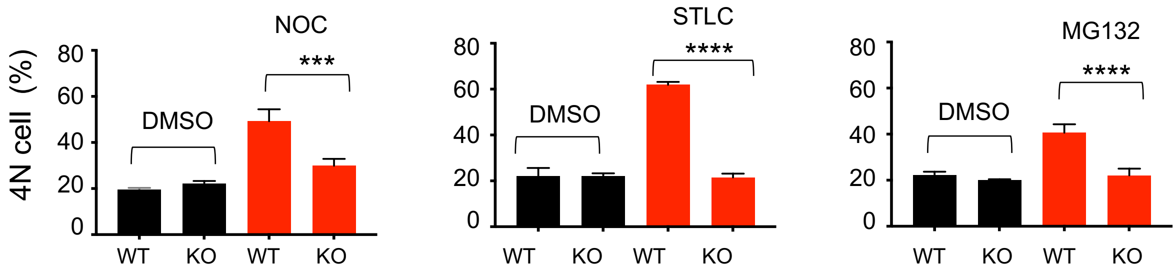


C Movies -Mitotic progression

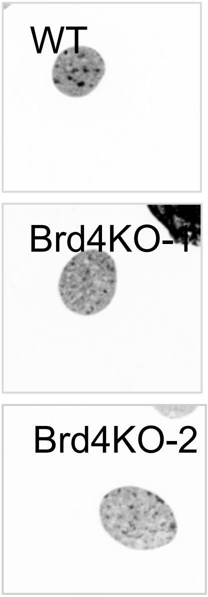


D Mitotic progression of WT and Brd4KO cells

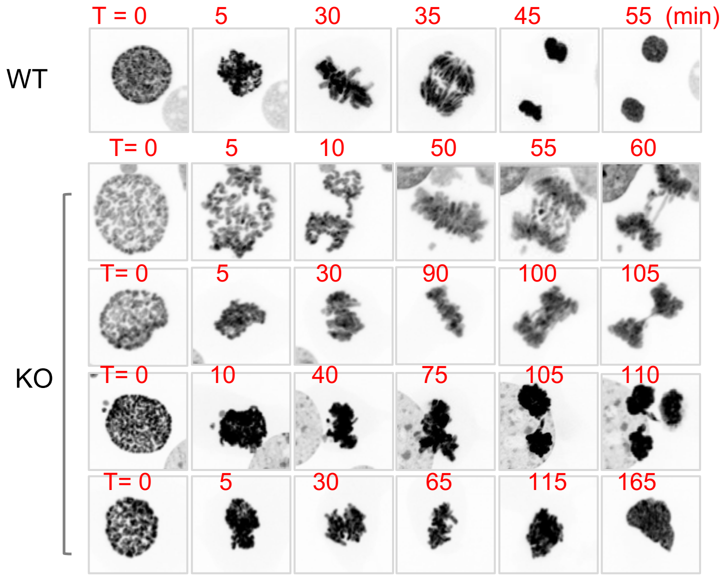

**Supplementary information, Figure S5**

**Effects of drugs that arrest mitotic progression**

A. Asynchronously growing WT and Brd4KO cells were treated with Nocodazole (100 mg/ml), or STLC (5 μM), or MG132 (5 μM) for 8h, stained with propidium iodide, and DNA contents were analyzed by flow cytometry. The numbers represent the percentage of the cells.

B. Quantification of 4N cells. Values represent the means of three independent experiments +/- S.D. Significance was calculated using unpaired t test (*** P<0.0001; ** P<0.001).

C. Live cell images of WT and Brd4KO cells expressing H3.1-GFP.

D. Real time imaging of one WT (Top) and four Brd4KO cells (bottom) undergoing mitosis. Note that the WT cells completed mitosis to produce two daughter cells in 55 min. Some Brd4KO cells underwent unequal chromosomal segregation (middle three) or disintegrated without producing progeny (bottom, See also Fig. 5F as well).

**Supplementary information, Figure S6**

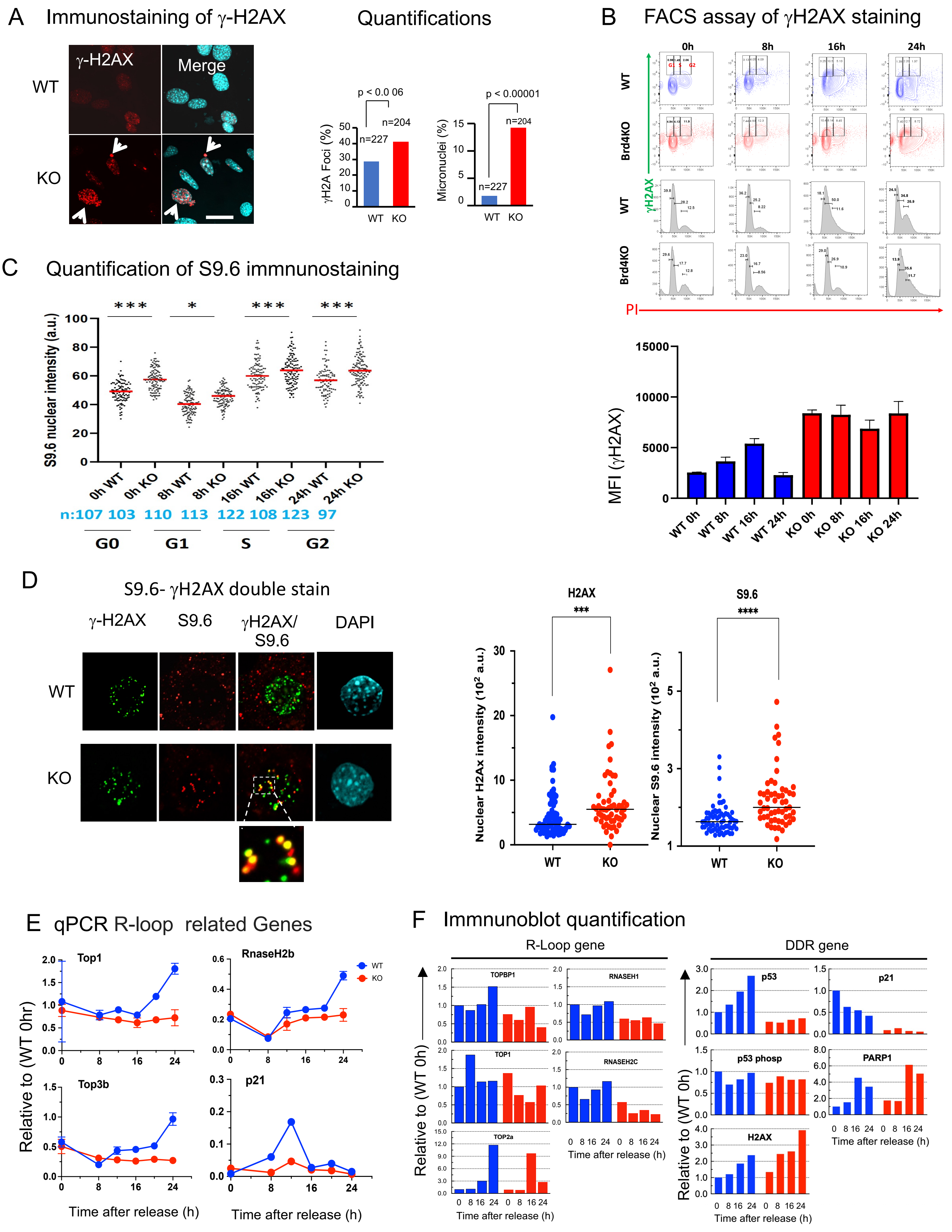

### G Genome view of R-loop and DDR genes

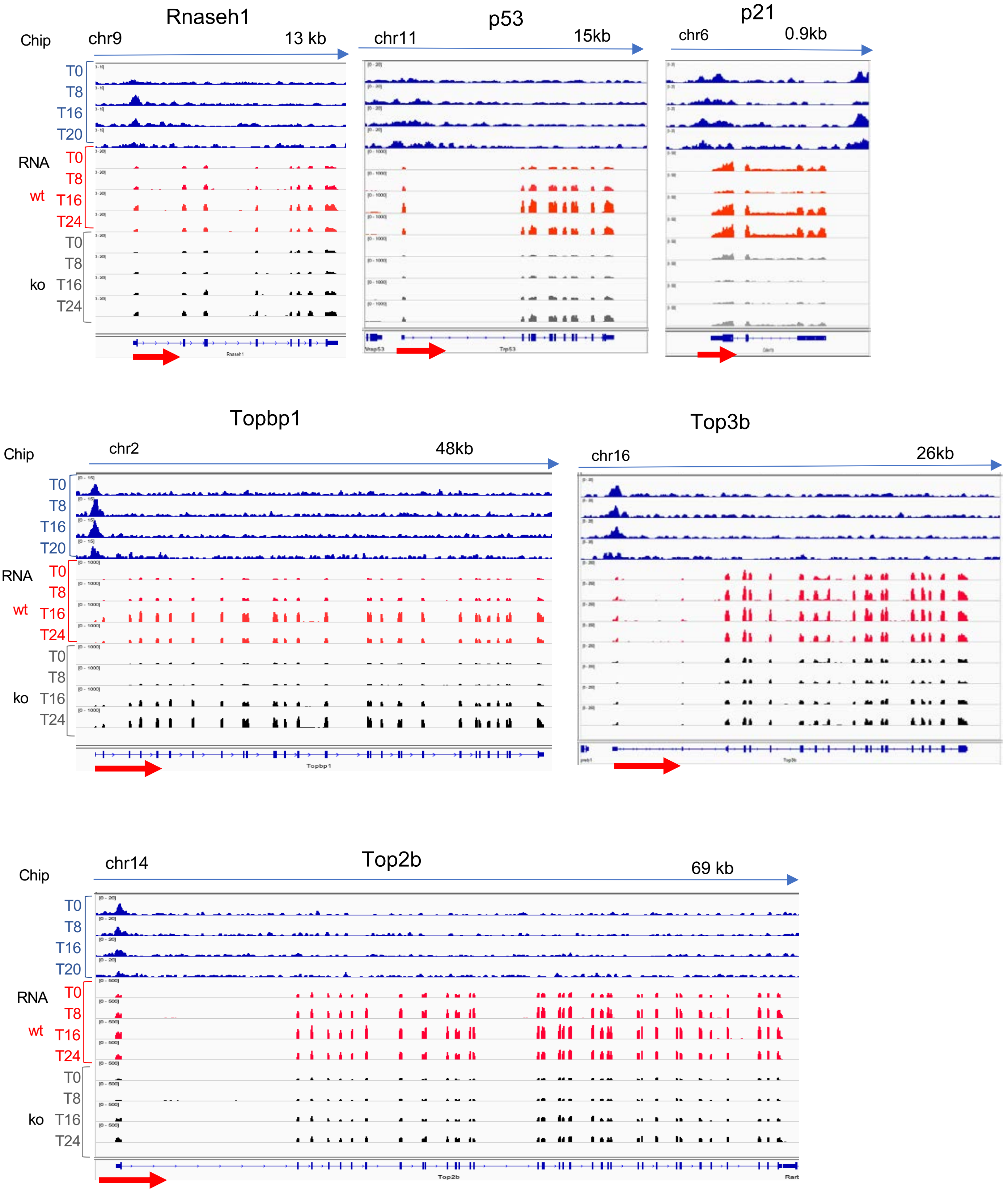

**Supplementary information, Figure S6**

**Real time imaging of mitotic progression**

A. WT and Brd4KO cells were immuno-stained with γH2AX antibody, and counterstained with Hoechst 33342 for DNA. Arrow indicates cells positive with γH2AX foci (left). Quantification of cells with γH2AX positive nuclei and micronuclei particles (right).

B. FACS profiles of γH2AX positive cells. WT and Brd4KO cells were synchronized by serum starvation, released, and allowed to proceed for indicated times, then stained with PI (X axis) and γH2AX antibody (Y axis). In upper panels, γH2AX positive cell populations are marked by a rectangle. Lower panels are histogram representation of corresponding cells. Bottom panel represents mean fluorescent intensity (MFI) of γH2AX signals. Significance was assessed by the unpaired t test (*** $P<0.0001$; ** $P<0.001$ Bottom).

C. Quantification of nuclear S9.6 intensity was performed using ImageJ (Figure 6E) (*$P <0.05$ to ***$P < 0.001$), significance of IF intensity was assessed using two-tailed unpaired t test.

D. Randomly growing WT and Brd4KO cells were stained with antibodies for γH2AX (green) or S9.6 (red) or both, and counterstained with DAPI (right). Enlarged image of γH2AX and S9.6 foci in Brd4KO cells showing colocalization of the two (bottom). Quantification of γH2AX and nuclear S9.6 intensity per cell (right). Significance assessed using Mann-Whitney test at right (γH2AX ***$P= 0.001$; S9.6 ****$P < 0.0001$).

E. qRT-PCR confirmation of R-Loop related genes in synchronized WT and Brd4KO cells. mRNA expression was normalized to Hprt. Values represents the average of three independent experiments.

F. Quantification of immunoblots Figure 6F. β–ACTIN was used as a loading control.

G. IGV profiles of BRD4 occupancy (top) and RNA-seq peaks (middle and bottom) for R-loop and DNA damage genes in WT and Brd4KO cells. Gene names and the exon-intron organization are shown below.

H. BRD4 integrates R-loop and DNA damage control into cell cycle progression

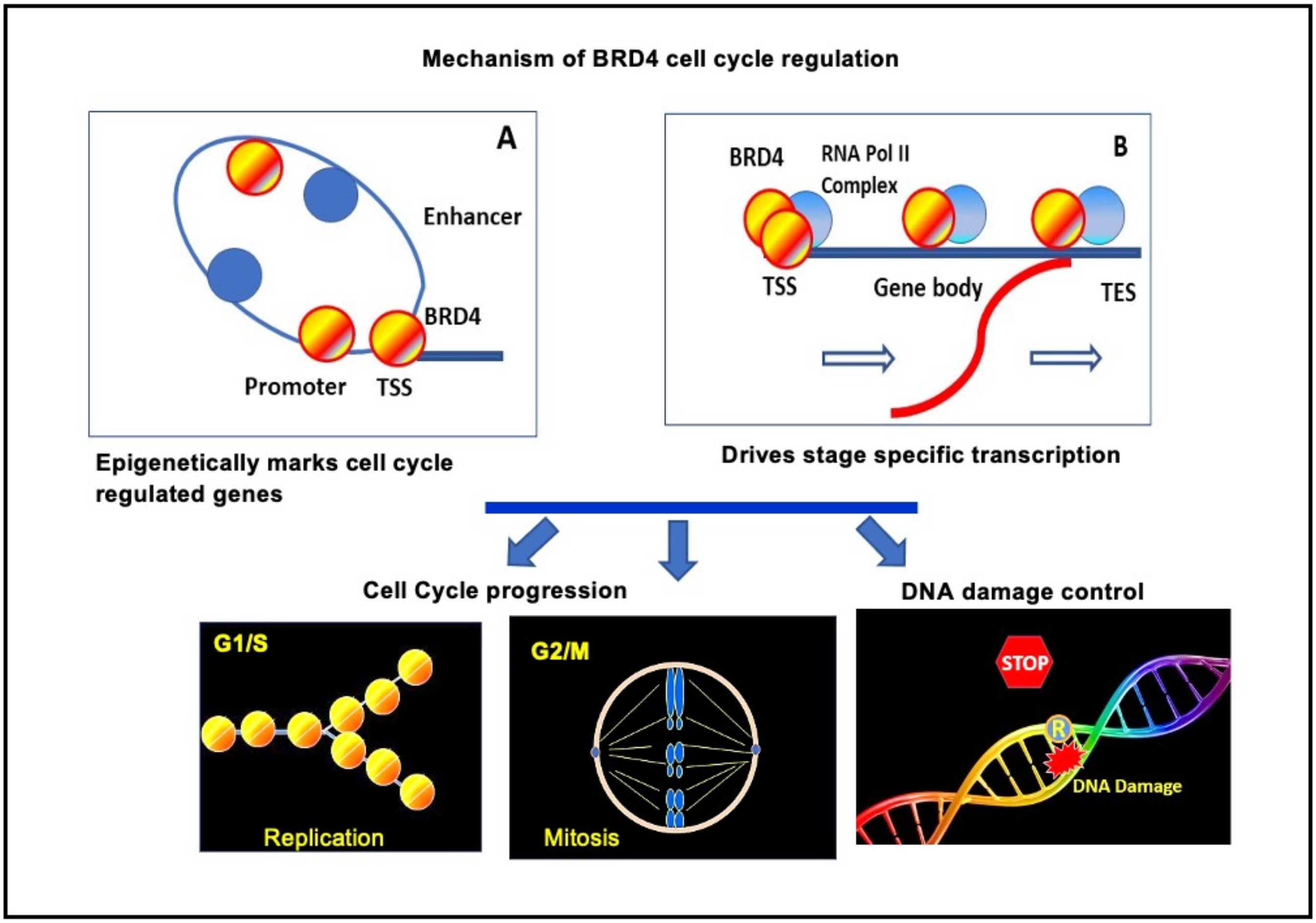


Mechanism of BRD4 action in cell growth

- BRD4 occupies many cell cycle genes that drives S phase passage and mitotic cell division continuously throughout cell cycle stages, providing stable epigenetic marks (left). This process is coupled with stable marking of genes regulating R-loop formation and DNA damage response (DDR).
- BRD4 drives timely transcription of cell cycle genes and those that regulate R-loop formation and DDR (right).
- This study demonstrates that DDR is an integral part of proliferation both normal and cancer cells.
